# Inhibiting B-cell-mediated Immunosuppression to Enhance the Immunotherapy Efficacy in Hepatocellular Carcinoma

**DOI:** 10.1101/2025.03.04.641494

**Authors:** Xin Liu, Zelong Liu, Tatsuya Kobayashi, Pin-Ji Lei, Yue Shi, Dandan Yuan, Jianguo Wang, Min Li, Aya Matsui, Kassiana Mafra, Peigen Huang, Ming Kuang, Lloyd Bod, Dan G. Duda

## Abstract

**Background:** Immunotherapy is efficacious in hepatocellular carcinoma (HCC), but the benefits are limited to a minority of patients. Most HCC patients show resistance to immune checkpoint blockade (ICB). Agonists of the stimulator of interferon genes (STING), potent immune stimulators, showed limited effectiveness. Using preclinical models, we studied the mechanisms of resistance to ICB and STING agonism.

**Methods:** Murine HCA-1 and RIL-175 HCCs were orthotopically grown in mice with underlying liver fibrosis, to mimic the presentation of human HCC. Established tumors were treated with a STING agonist (BMS-986301) or anti-PD1 ICB, and mice were followed to evaluate safety and efficacy, as well as the mechanisms of treatment resistance by RNA sequencing, flow cytometry, and immunofluorescence, B-cell depletion and T-cell immunoglobulin and mucin domain 1 (TIM-1) ICB.

**Results:** Unbiased analyses of transcriptomic data from murine HCC tissues from ICB-treated mice showed an increased abundance of intratumoral CD8^+^ T cells and B cells. STING agonism alone showed efficacy in the ICB-responsive RIL-175 HCC model but more limited efficacy in the ICB-resistant HCA-1 model. STING agonism increased circulating IL-10 and intratumoral infiltration by B-cells, including TIM-1^+^ B cells, and promoted the formation of tertiary lymphoid structure (TLS)-like structures, especially in the peritumoral areas. Strikingly, adding B cell depletion to ICB or STING agonism treatment significantly increased survival. Interestingly, unlike ICB, STING agonism also had a pronounced anti-metastatic activity. In addition, the combination of STING agonism and TIM-1 blockade augmented B cell differentiation and antigen presentation *in vitro* and improved the anti-tumor effects in murine HCC *in vivo*. This approach decreased the number of TIM-1^+^ B cells in the tumor and shifted B cells to higher expression of CD86 and MHC class II, enhancing the antigen presentation capability and further boosting the antitumor efficacy of CD8^+^ cytotoxic T cells.

**Conclusion:** Our findings demonstrate that B cells are associated with ICB- and STING-mediated therapy resistance, and that depleting B-cells or targeting TIM-1 enhances both innate and acquired therapeutic efficacy in HCC.

## Introduction

Hepatocellular carcinoma (HCC) is one of the most common malignancies and a leading cause of cancer-related mortality worldwide, representing a major global healthcare challenge with increasing incidence and mortality (1–3). The majority of HCC development occurs in patients with underlying liver disease, mostly because of hepatitis B or C virus infection, while non-alcoholic steatohepatitis associated with metabolic syndrome or diabetes mellitus is becoming a dominant risk factor in Western countries (4,5). Although surgical and locoregional treatments are becoming more widely available globally, it is estimated that approximately 50–60% of patients with HCC will eventually receive systemic therapies (1,6). Systemic therapies have been the mainstay treatment of advanced HCC for almost two decades (7). The successes of the phase III trials of combination therapy based on immune checkpoint blockade (ICB) (anti-PD-L1 with anti-VEGF in the IMbrave150 study and anti-PD-L1 with anti-CTLA-4 in the HIMALAYA study) have transformed in systemic therapy for HCC (8,9). Despite this progress, more than 70% of the patients with advanced HCC do not respond to current ICBs and most suffer from disease progression. Enhancing immunotherapy approaches by targeting the immunosuppressive tumor microenvironment (TME) of HCC remains an urgent need.

As key components of adaptive immunity, T lymphocytes play a well-established role in tumor responses to ICB (10). However, the contribution of B lymphocytes to this process remains less understood, with most insights emerging only in the past decade (11). Within the TME, B cells can exhibit a broad spectrum of cell states and mediate innate and adaptive immune responses (12,13). Their antigen-presenting ability enables them to generate co-stimulatory or co-inhibitory signals and release cytokines and chemokines, modulating the behavior of other cells, such as effector T cells (12,13). Tumor-infiltrating B cells may exert both pro-tumor and anti-tumor effects depending on their phenotypes and TME composition. The formation of tertiary lymphoid structures (TLS) with B-cell follicles in cancer tissues, including HCC, indicates the crucial role of B cells and TLS in mediating anti-tumor immunity (14–17). Conversely, regulatory B cells (Bregs) inhibit immune responses to maintain immune homeostasis and promote tumor progression (18,19). Reports suggest that certain B-cell subsets and antibody specificities could contribute to cancer relapse and metastasis (20,21). Our recent study uncovered a B-cell population that expands during tumor progression, marked by the surface receptor T-cell immunoglobulin and mucin domain 1 (TIM-1) and other T-cell checkpoint receptors, indicating that TIM-1 marks a subset of activated B cells expressing co-inhibitory molecules and IL-10. This B cell subset significantly impairs anti-tumor T cell responses in multiple murine cancer models (22). These findings underscore the need to further elucidate the roles of specific B-cell subpopulations and cell states in immunotherapy resistance in HCC.

A pathway identified as critical for the innate immune system and anti-tumor immunity is the cyclic GMP-AMP synthase (cGAS)-stimulator of interferon genes (STING) signaling (23,24). STING proteins, located in the endoplasmic reticulum, facilitate innate immune signaling by inducing the expression of type I interferons (IFNs) and pro-inflammatory cytokines upon sensing cytosolic double-strand DNA (25–27). Mechanistically, the binding of STING agonist to STING recruits TANK-binding kinase 1 and interferon regulatory factor 3, leading to the production of type I IFNs and pro-inflammatory cytokines, which results in the maturation, migration, and activation of dendritic cells, T cells, and natural killer (NK) cells (28–31). STING agonist agents are actively being evaluated in pre-clinical and clinical studies as enhancers of anti-tumor immune responses. However, while modulating STING has shown promise against primary and metastatic cancers in pre-clinical studies, STING agonists have demonstrated limited anti-tumor efficacy in the clinical trials conducted so far, even when combined with PD-1/PD-L1 ICB (32–34). This underscores the need to counteract immunosuppressive factors within the TME when employing STING agonists. Notably, STING activation has been shown to induce regulatory B cells that impair NK cell function in pancreatic cancer (35), suggesting a potential immunosuppressive role for Bregs in limiting STING-mediated anti-tumor immunity. However, the precise mechanisms remain incompletely understood, and these mechanisms are entirely unknown in HCC, restricting the development of effective combination strategies despite the promising activity of STING agonists in preclinical models (36,37).

Since the discovery of STING, a range of natural and synthetic STING agonists have undergone evaluation in pre-clinical and clinical settings for different tumor types (38). The notable pre-clinical anti-tumor effects of STING agonists have led to the development of multiple pharmacologic classes of agents, including cyclic dinucleotides, non-cyclic dinucleotides, bacterial vectors, and other unique STING agonists (28,38,39). Currently, several STING agonists, such as ADU-S100/MIW815, E7766, and GSK3745417, have been approved for clinical trials in treating solid tumors or lymphoma (28,40). However, some have been terminated because no substantial anti-tumor activity was observed in humans. The challenge of delivering STING agonists into the cytosol has led to the predominant use of intratumoral injections for administration in clinical trials. This approach enables precise tumor targeting, achieving high local concentrations with reduced systemic distribution and toxicity. Nonetheless, it brings specific challenges for further application in HCC treatment, such as the risks of bleeding and needle tract implantation (41).

BMS-986301 is a novel STING agonist undergoing clinical trials as a systemic treatment using intramuscular injections, either alone or in combination with nivolumab and ipilimumab, for advanced solid cancers resistant to checkpoint inhibitor therapy (NCT03956680). Preliminary results have demonstrated that BMS-986301 monotherapy achieved over 90% complete regression in murine models of colorectal cancer, exhibiting lower toxicity towards CD8^+^ T cells and less inhibition of their proliferation compared to ADU-S100.

In this study, we evaluated the efficacy of STING agonism in murine models of HCC with underlying liver damage, which mimics the presentation of human disease. Our findings illuminate the pivotal role of B cells in STING agonist-based HCC treatment and offer insights into overcoming resistance to such therapies. By harnessing B-cell-mediated immunity, particularly by identifying specific targets on B cells, we can amplify the efficacy of existing HCC immunotherapies. This approach also opens avenues for treating tumors previously unresponsive to treatment with STING agonists alone, providing a promising strategy for enhancing HCC management.

## Methods

### Cells and culture condition

We used 2 murine HCC cell lines: HCA-1 from C3H mice, established in our laboratory (42,43), and RIL-175 (a *p53*/*Hras* mutant HCC cell line from C57Bl/6 mice, a kind gift from Dr. Tim Greten, NIH) (44). HCA-1 was maintained in Dulbecco’s Modified Essential Medium (DMEM) (ThermoFisher, USA) with 10% fetal bovine serum (FBS) (Hyclone, SH30071.03) and 1% penicillin-streptomycin (Gibco #15070063) in 5% CO_2_ at 37℃. RIL-175 was maintained in DMEM with 20% FBS and 1% penicillin-streptomycin in 5% CO_2_ at 37℃. All cells used for experiments were passaged less than 5 times and were authenticated before in vivo use. Mycoplasma contamination was routinely performed before in vivo studies for all cell lines using MycoAlert Mycoplasma Detection Kit (Lonza #LT07-318). No genetic manipulations were performed for the cells used in this study.

### Animal studies

Animal experiments were performed in the animal facility of Massachusetts General Hospital under specific pathogen-free conditions. All animal experiments were performed under the Institutional Animal Care and Use Committee (IACUC) at Massachusetts General Hospital-approved protocol (2020N000023). Studies complied with all guidelines outlined regarding animal research in the IACUC Policies and Guidance of MGH Research Institute.

### Orthotopic HCC mouse models

In the therapeutic studies, orthotopic HCCs were induced by intrahepatic injection of HCA-1 cells in syngeneic C3H mice, while RIL-175 cells were implanted in syngeneic C57Bl/6 mice. The mice were purchased from the MGH Center for Comparative Medicine. Six-to-8-week-old male mice were used for experiments. HCC model under liver damage was performed as described previously (43). To induce liver damage, 100μl of 20% carbon tetrachloride (CCl4) (Sigma-Aldrich #289116) was administrated orally for 6 weeks. After one week of recovery, the suspensions of the murine HCC cells mixed with Matrigel (Corning #354234) in a 1:1 volume ratio were injected into the subcapsular region of the liver parenchyma using 0.5ml syringes with 28-gauge needles. To prevent leakage of tumor cells from the injection sites, the injection volume was controlled to 10μl (10^6^ cells in 10μl per mouse). In addition, a steady and slow injection was performed to minimize leakage of the injected cell suspension further. After withdrawing the needle, the injection site was covered with Surgifoam (Ethicon #1972) for 5 minutes to reduce bleeding and potential backflow. Treatments were initiated in mice with established tumors when the tumors reached 5mm in diameter, measured by high-frequency ultrasound imaging. Tumor growth and treatment response were also monitored by ultrasound imaging.

### Imaging of orthotopic HCC

Tumor growth and treatment response were monitored by high-frequency ultrasound imaging. For the longitudinal evaluation of tumor growth, we used an ultrasound device (Vevo 2100, VisualSonics) equipped with specific probes for small-animal imaging weekly. Imaging to assess tumor growth longitudinally was conducted noninvasively under isoflurane anesthesia. The ultrasound measurement was discontinued upon the demise of over 50% of the mice in a treatment group and the health status of the mice to recover from the anesthesia.

### Treatments

BMS provided STING agonist BMS-986301. Mouse anti-CD19 (clone 1D3), anti-B220 (clone RA3.3A1/6.1), anti-TIM-1 (clone 3B3), anti-PD1 (clone RMP1-14), and anti-VEGFR2 (clone DC101) were purchased from BioXCell (Lebanon, NH). The STING agonist was administered by intramuscular (i.m.) injection (2mg/kg, weekly, 2-3 doses). Anti-CD19 and anti-B220 depleting antibodies (10 mg/kg, every 5 days for 3 weeks), anti-TIM-1 blocking antibodies antibody (10 mg/kg, every 5 days for 20 days), and anti-PD1 and anti-VEGFR2 blocking antibodies were administered by intraperitoneal (i.p.) injections (anti-PD1: 10mg/kg, anti-VEGFR2: 20mg/kg, every 3 days for 21 days). Corresponding isotypes of IgG were administered i.p. at the same frequency as the other antibodies. B-cell depletion was validated by flow cytometry analysis of peripheral blood mononuclear cells collected 5 days after the last antibody dose.

### Immunofluorescence

Six-µm-thick frozen sections of murine HCC tissue were prepared for immunofluorescence (IF). We used an anti-CD31 antibody (Millipore, clone 2G8) to identify endothelial cells, an anti-α-SMA antibody (Sigma, clone 1A4) to identify perivascular cells, anti-CD3 (Abcam #ab135372), anti-CD8 (CST, #98941) and anti-CD4 (Abcam, #ab288724) antibodies for T cells staining, and anti-CD19 (Abcam #ab245235) and anti-B220 (R&D #MAB1217) antibodies for B cells staining. All secondary antibodies were purchased from Jackson ImmunoResearch (West Grove, PA, USA). Frozen sections from OCT-embedded tissue blocks were washed with PBS and treated with normal donkey serum (Jackson ImmunoResearch #017000121) for blocking. Primary antibodies were applied overnight at 4℃, followed by the reaction with corresponding secondary antibodies for 1 hour at room temperature. Analysis was performed in random fields in the tumor tissues under ×400 magnification using a laser-scanning confocal microscope (Olympus, FV-1000). Whole-slide scanning was conducted using Zeiss Axio Scan Z1. Data were analyzed using ImageJ (US NIH) and QuPath software (45).

### Lung preparation

Complete lungs were dissected at the hilum from the pulmonary trunk of the heart and immediately immersed in Bouin’s fluid (Electron Microscopy Sciences, #15990-01) for 24 hours for the long-term experiment. Lung metastatic burden was assessed by enumerating metastatic nodules on the surface of the lung.

### H&E staining

Five-μm-thick sections of murine HCA-1 lung metastatic tissues were deparaffinized in xylene for 5 minutes x 2 times, rehydrated using a graded alcohol series, and placed in a citrate buffer at 97°C for 20 minutes for antigen retrieval. After hydration, hematoxylin was applied for 2 minutes to stain cell nuclei, followed by a brief rinse in water. Eosin (Sigma-Aldrich #102439) was applied for 1 minute to stain the cytoplasm. After mounting, slides were taken images using a bright-field microscope (Olympus, BX40) under ×40 magnification by a Canon camera. Data were analyzed using QuPath software (45). 3 regions were randomly selected for each sample for statistical analysis.

### RNA sequencing

Total RNA was extracted from the HCC tissues using Qiagen kits. RNA sequencing (RNA-seq) was performed on Illumina Novaseq at the MIT BioMicro Center (Cambridge, MA). FastQC was performed for the quality control of RNA-seq raw data. After quality control, the low-quality bases and adaptors contamination was removed by Cutadapt. The quality of clean yield data was examined again by FastQC software. Next, the clean data were aligned to the mouse reference genome mm39 by STAR. After data mapping, samtools and featureCounts were used to count the number of reads aligned to the gene features. DESeq2 identified the differentially expressed genes. Differentially expressed genes were annotated using REACTOME databases, and the cell type enrichment analysis was performed using the xCell package (46). The network plot of enriched pathways was performed by the clusterProfiler package. Single-sample gene set enrichment analysis (ssGSEA) was used to calculate the Breg score based on the markers (47). TCGA data analyses were performed with log-rank Mantel-Cox test using web server GEPIA2 (48), based on TCGA liver hepatocellular carcinoma (TCGA-LIHC) data.

### Flow cytometry analysis

Harvested cells were washed with the buffer and stained with the cell surface antibodies. Anti-mouse CD16/32 antibody (clone 93, Biolegend, San Diego, California, USA) was added for FcR blockade and incubated for 5 minutes at room temperature; 7-amino-actinomycin D was added for live/dead staining. After another washing step, antibodies for cell phenotyping were added, and cells were incubated for 30 minutes at room temperature. The monoclonal antibodies used for flow cytometry analysis were specific for CD45 (BioLegend, 30-F11), CD3 (BioLegend, 17A2), and CD19 (BioLegend, 1D3/CD19).

### Cytokine analysis

Mouse plasma samples were assayed in duplicate using the MSD V-PLEX proinflammatory panel 1 mouse kit, a highly sensitive multiplex enzyme-linked immunosorbent assay (ELISA) for quantitatively measuring 10 cytokines: IFN-γ, interleukin (IL)-1β, IL-2, IL-4, IL-5, IL-6, IL-10, IL-12p70, CXCL1, and tumor necrosis factor (TNF)-α from a single small sample volume (25μl) using electrochemiluminescence-based detection (MesoScale Discovery, Gaithersburg, MD).

### Statistical analysis

Mann-Whitney U test was utilized to compare two groups with quantitative variables. When the experimental cohort includes more than two groups, including quantitative variables, one-way ANOVA with Tukey’s multiple comparisons test was applied unless specified in the figure legends. The Kaplan-Meier method generated survival curves underlying the Log-Rank test and Cox proportional hazard model. The hazard ratio (HR) and 95% CI were calculated for statistical survival analyses for murine models. All analyses were performed using GraphPad Prism 9 (GraphPad Software, MA, USA), and data were presented as mean values ± SD. The significant difference between experimental groups was determined when p-values were less than 0.05.

### Data availability statement

Data are available on reasonable request from the corresponding author DGD. Reagents are available via materials transfer agreements (MTAs).

## Results

### ICB immunotherapy drives B-cell infiltration in murine HCC

To objectively assess changes in intratumoral immune cell populations following ICB, we integrated seven bulk RNA-seq datasets of murine liver cancer from our previous studies (49–53). Using anti-PD-1 antibody as the primary treatment, principal component analysis (PCA) revealed significant transcriptional shifts in response to ICB (**Fig. 1A, B**). Immune deconvolution revealed a significant increase in two major immune populations following treatment: CD8^+^ T cells, as expected and previously reported (51,52), and more notably, B cells among non-tumor cell populations (**Fig. 1C, D**). Multiple immune profiling approaches consistently identified this expansion, with large effect sizes quantified using Cohen’s d across various methods (**Fig. 1C**). Notably, the increase in B cells post-ICB treatment was further validated through multiple deconvolution techniques, underscoring the robustness of these findings (**Fig. 1E**). As ICB immunotherapy triggers adaptive immune responses, our findings indicate that the increased B cell infiltration is associated with the activation of adaptive immunity, highlighting its potential role in the response to immunotherapy.

**Figure 1.**
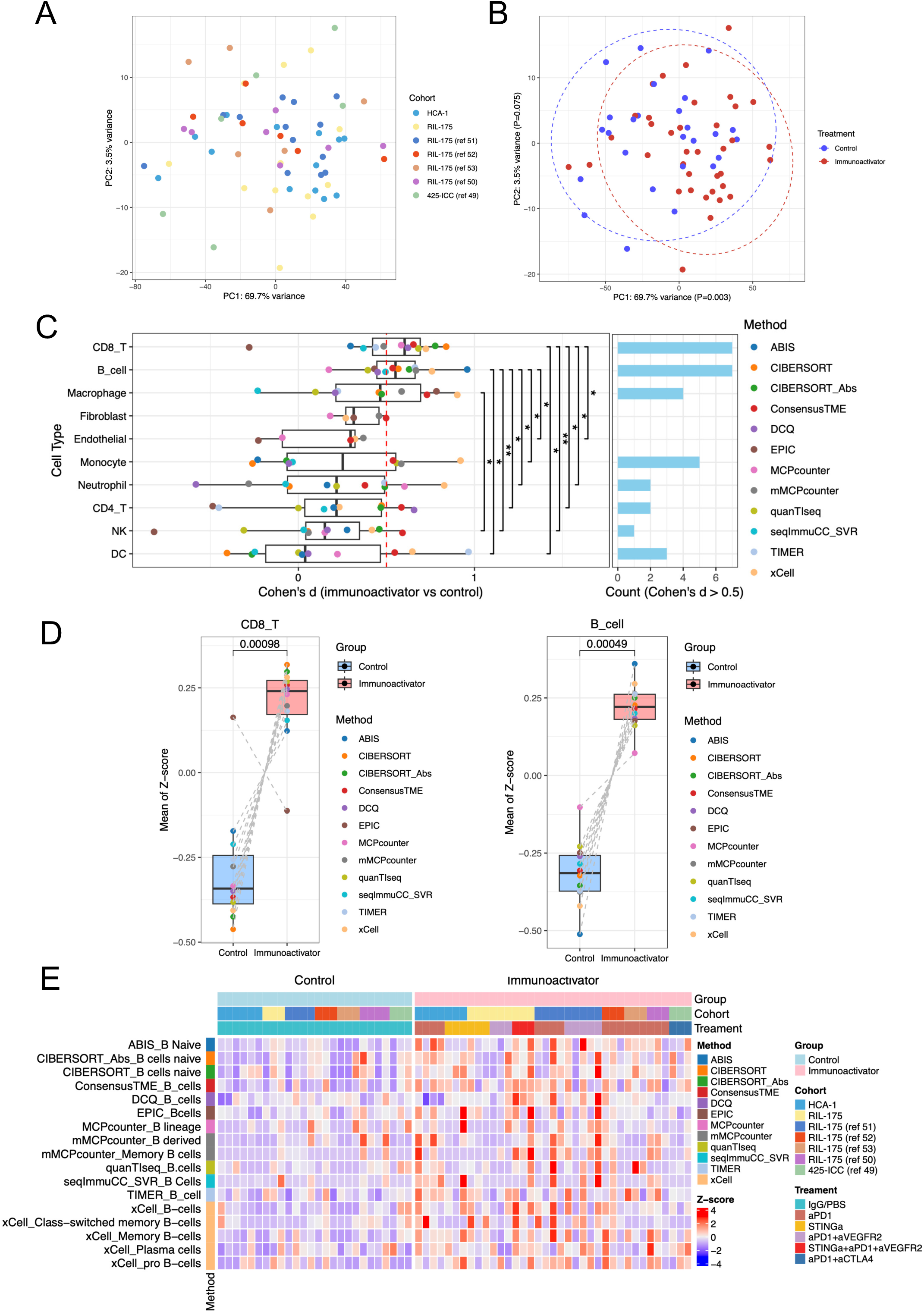
ICB immunotherapy increased B-cell infiltration. (**A**) PCA plot of RNAseq data from 5 published studies and 2 immunotherapy-treated cohorts (49–53). (**B**) PCA plot of RNAseq data showing the difference between immunoactivator (n=37) and control (n=26) groups. PC1: p=0.003 (t-test). (**C**) Significant increases in CD8^+^ T and B cells were detected using multiple immune deconvolution methods. Cohen’s d statistic was used to quantify the effect size of proportion change of each predicted cell subset in the ICB and control groups. Cohen’s d classified effect sizes as small (d = 0.2), medium (d = 0.5), and large (d ≥ 0.8). (**D**) The mean of the Z-score of the deconvolution score of each group was predicted by 11 methods. P<0.05 from paired Wilcoxon test. (**E**) Deconvolution results of B cells in TME of liver cancer by 11 methods.

### STING agonist treatment enhances B-cell infiltration in HCC

Next, to understand the alterations in the TME following the activation of innate immunity, we investigated the effects of enhancing anti-tumor immunity by using a STING agonist (BMS-986301) in the ICB-responsive RIL-175 murine HCC model in C57Bl/6 mice. First, to determine its optimal dosing, mice with established tumors and underlying liver damage received 2 i.m. injections of STING agonist at varying doses: 0.67, 1.33, 2.00, 2.67, and 3.33 mg/kg, administered weekly. We monitored the changes in body weight as a measure of toxicity and tumor growth delay by ultrasound imaging. All doses led to body weight loss in some mice, notably one week after the treatment, yet all mice recovered to their normal weight shortly thereafter (**Fig. S1A**). Doses higher than 2 mg/kg were associated with improved therapeutic outcomes, manifesting as delayed tumor growth and extended survival compared to lower doses (**Fig. S1B**, **C**). Given the potential risk of high doses of the STING agonist inducing T cell apoptosis and adverse effects (54), we selected the 2 mg/kg weekly dose that showed anti-HCC activity without limiting toxicity in mice with liver damage for further testing of the efficacy and safety of the treatment.

Next, we tested the efficacy of the STING agonist in a highly metastatic and ICB-resistant murine HCC model (orthotopic HCA-1 grafted in C3H mice) (51). Weekly administration of the STING agonist for two doses of 2 mg/kg induced a transient growth delay but did not increase median OS in this model (**Figs. 2A** and **S1D**). Flow cytometric analysis of the tumor tissues from a separate time-matched cohort showed that the proportion of tumor-infiltrating B cells was significantly higher after STING agonism (**Fig. 2B**). We conducted bulk RNA-sequencing (RNA-seq) analysis on tumor tissue samples collected on day 10 post-treatment to investigate the alterations within the TME that may mediate immunosuppression. The analysis revealed an upregulation of B-cell-related pathways in the STING agonist-treated group compared to the control group (**Fig. S2A**). These upregulated pathways included FCGR3A-mediated IL10 synthesis, CD22-mediated BCR regulation, and signaling by the B cell receptor (**Fig. S2B, C**), motivating us to focus on the enrichment of B cells. Using the bioinformatic tool xCell, which is designed to perform cell type enrichment analysis from gene expression signature (46), we found that the enrichment score of total B cells increased after STING agonist treatment compared to the control group, with memory B cells showing the most increase (**Fig. 2C, D**). The increased B-cell infiltration was also confirmed by IF, which showed a significant increase in B-cell proportion after STING agonist treatment (p=0.0005) (**Fig. 2E, F**).

**Figure 2.**
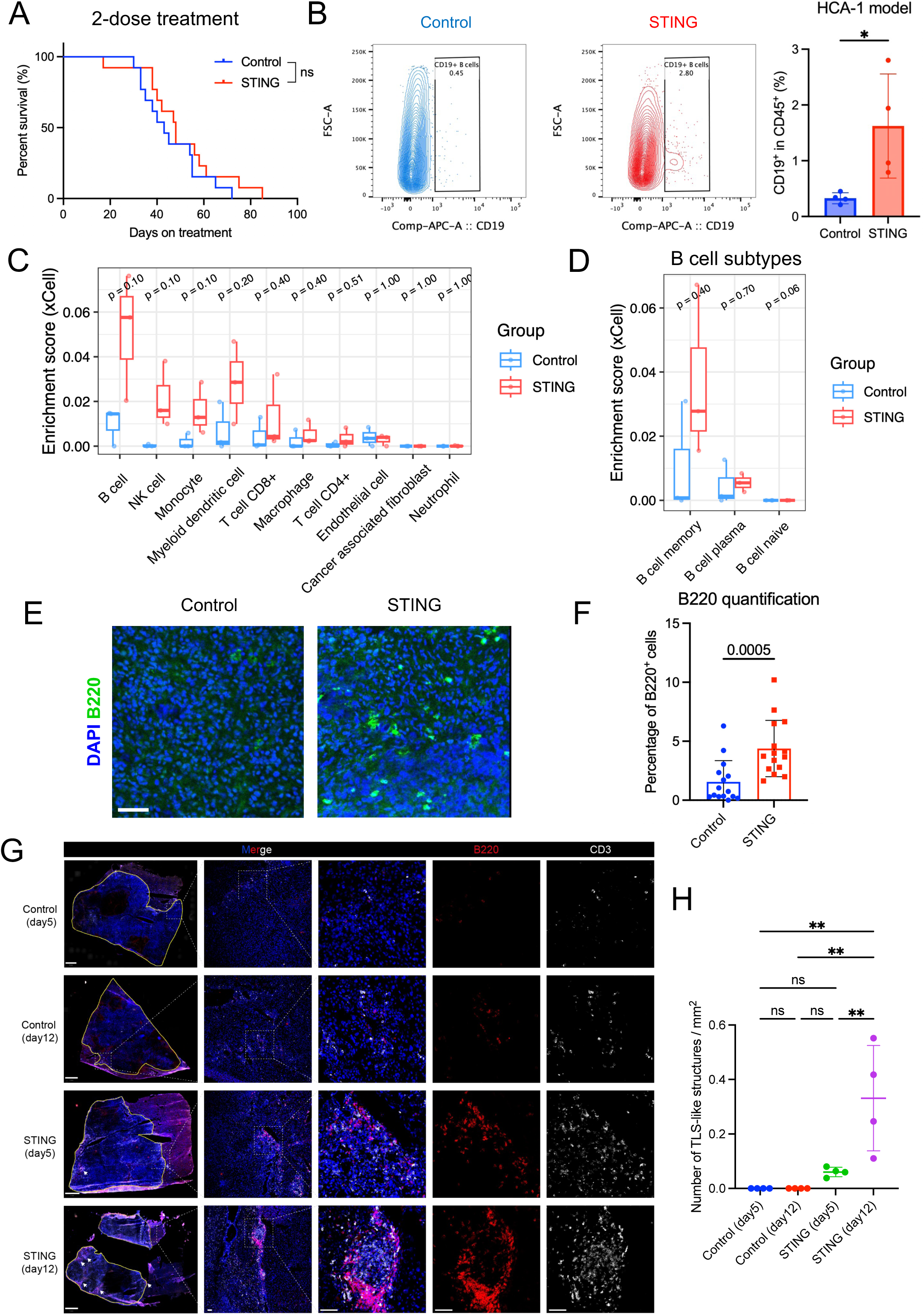
B cells are enriched after STING agonist treatment in orthotopic HCA-1 and RIL-175 murine HCCs. (**A**) Overall survival of HCC-bearing mice after weekly administration of STING agonist treatment or control for two doses. Log-rank test, n=13/group. (**B**) Flow cytometry analysis of intratumor CD19^+^ B cells in both groups. Comparison of the proportion of CD19^+^ cells in CD45^+^ cells from the HCA-1 orthotopic HCC tumors between STING agonist-treated and control groups (n=4/group). Statistical significance was calculated by the Mann-Whitney U test. (**C**) The enrichment score of major cell types in the HCA-1 tumor samples from STING agonist and control groups was calculated by xCell. (**D**) Enrichment score calculated by xCell of subtypes of B cells between STING agonist-treated and control groups. (**E**) Representative immunofluorescence (IF) for the B-cell marker B220 among the two groups (scale bar, 50μm). (**F**) Comparison of percentage of B220^+^ cells of tumor samples collected from HCA-1 murine HCC, showing significantly higher infiltration of B cells after STING agonist treatment (Mann-Whitney U test). (**G**) Representative IF results for tissues collected from control and STING-agonism-treated mice on days 5 and 12. Scale bar=1mm on the left column, other scale bar=50μm. Tumor regions were delineated by yellow lines. (**H**) Statistical comparison of the number of TLS-like structures per mm^2^. **, P < 0.01, statistical significance was calculated by one-way ANOVA with Tukey’s test.

Next, we tested the effects of STING agonism in the orthotopic RIL-175 murine HCC model in C57Bl/6 mice with liver damage, which responds to anti-PD1 therapy (51). We found that treatment with the STING agonist alone increased the number of B-cell aggregates compared to the control group after just one dose (**Fig. S3A, B**), indicating a rapid functional reprogramming of B-cells after the treatment. We performed staining using CD3 and B220, identifying TLS-like structures in tumors from the STING agonist treatment group, particularly on day 12 (**Fig. 2G, H**). Most of these TLS-like structures were located in the peritumoral area, with very few observed within the intratumoral region (**Fig. S3C**). Moreover, we found that STING agonist treatment significantly increased the total and pericyte-covered microvessel density (**Fig. S4A, B**), and the intratumoral infiltration by CD4^+^ and CD8^+^ T-cells in the RIL-175 model (**Fig. S4C**). These data demonstrate that the STING agonist treatment promotes vascular normalization and enhances T-cell infiltration.

### B-cell depletion increases the therapeutic benefits of immunotherapy in an ICB-resistant murine HCC model

Given that increased B-cell infiltration during ICB treatment may indicate both pro-tumor and anti-tumor effects, we next performed B-cell depletion experiments in combination with ICB treatment. In the anti-PD1-resistant orthotopic HCA-1 murine HCC model, we tested the treatment efficacy of B cell depletion combined with dual PD1/VEGFR2 blockade (**Fig. 3A**). The combination of dual anti-PD1/VEGFR2 and B-cell depletion showed superior tumor growth delay (**Figs. 3B** and **S5A**), and significantly longer overall survival (OS) than other groups (**Fig. 3C**). No obvious adverse effects were observed as the body weight remained stable during the treatment period, and no differences in lung metastasis were found among the treatment groups (**Fig. S5B, C**). The efficiency of B-cell depletion was confirmed by flow cytometry analysis of blood samples collected on day 12 (**Fig. S5D, E**). These results indicate that the increase in intratumoral B cells after ICB immunotherapy has a predominantly immunosuppressive function in an immunotherapy-resistant murine HCC model.

**Figure 3.**
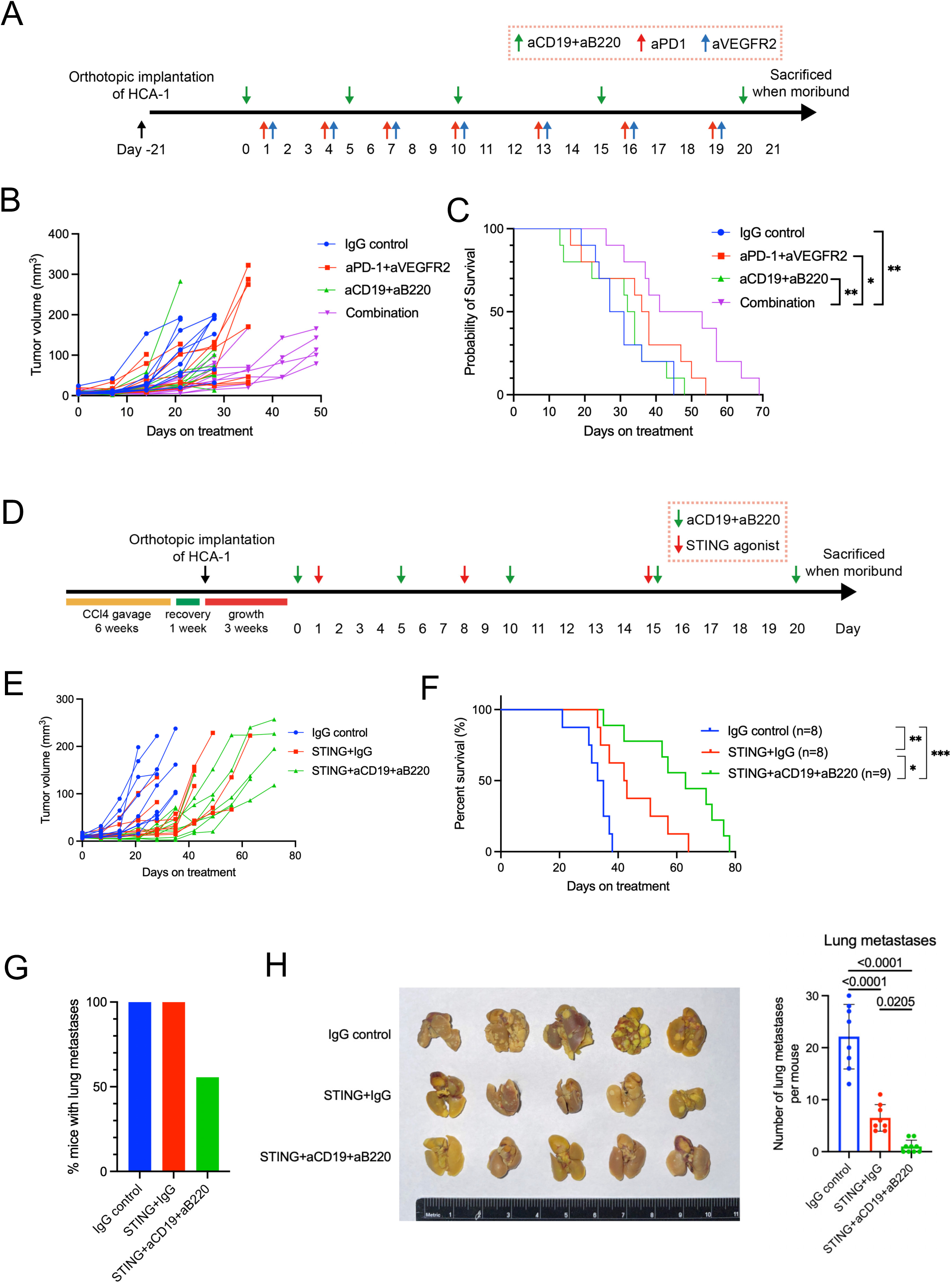
B cell depletion enhances the anti-tumor efficacy of immunotherapy and decreases metastasis in the murine HCA-1 HCC model. (**A**) Experimental design using the orthotopic HCA-1 murine HCC model. (**B**) Tumor growth kinetics after treatment: the combination of dual PD1/VEGFR2 blockade and B-cell depletion group induced tumor growth delay superior to other groups. (**C**) Overall survival of HCC-bearing mice after treatment: Combination of dual PD1/VEGFR2 blockade plus B cell depletion induced a significant survival benefit to other groups. Log-rank test, n=10/group. (**D**) Experimental design of B cell depletion with STING agonist treatment. (**E**) Tumor growth kinetics after treatment: the combination of STING agonist and B cell depletion group induced tumor growth delay superior to other groups. (**F**) Overall survival of HCC-bearing mice after treatment: STING agonist plus B cell depletion induced a significant survival benefit than STING alone or IgG control groups. Log-rank test, n=25. (**G**) STING agonism/B cell depletion reduced lung metastasis rates. (**H**) Representative photographs of lungs after fixation in Bouin’s solution and STING agonism/B cell depletion combined treatment significantly reduced lung metastasis in the HCA-1 murine HCC model. P values were calculated by one-way ANOVA with the Tukey multiple comparisons test.

To investigate the role of the increased B cell infiltration after STING agonist therapy, we conducted a survival study in mice with or without B-cell depletion by anti-CD19 and anti-B220 antibodies in HCA-1 murine HCC-bearing mice with underlying liver damage (**Fig. 3D**). To sustain the tumor growth delay effect seen with 2 doses of STING agonist, we administered a third weekly dose in this cohort. Mice with established orthotopic tumors were randomized to treatment with (**i**) IgG control, (**ii**) STING agonist with IgG, (**iii**) STING agonist with anti-CD19 and anti-B220 antibodies for B cell depletion. We found that STING agonist combined with B cell depletion effectively delayed tumor growth and significantly prolonged survival without limiting toxicities (STINGa + aCD19/aB220 vs STINGa + IgG, HR=0.38, p=0.02, log-rank test) (**Figs. 3E, F** and **S6A**). These data demonstrate that the increased B-cell infiltration has an immunosuppressive role and limits the survival benefit of STING agonism. The lungs are the most common site of HCC metastatic colonization, accounting for 51% of all extrahepatic metastases, one of the key factors affecting its prognosis (55–57). The HCA-1 model is highly prone to lung metastasis (51,58). Therefore, we also measured the lung metastases in the survival experiment and found lung metastasis incidence was the lowest in the STING agonist with B-cell depletion group (**Fig. 3G**).

When we evaluated the lung metastatic burden by enumerating metastatic nodules, we found it significantly reduced in the group that received a combination of STING agonist and anti-CD19/anti-B220 B-cell depletion than other groups (**Fig. 3H**). Moreover, pleural effusions were lowest in the STING agonist with the B-cell depletion group (**Fig. S6B**). However, ascites and peritoneal metastasis incidence were comparable between the treatment groups at the terminal endpoint (**Fig. S6C**, **D**). H&E staining results showed that the STING agonist potently inhibited metastasis, and the addition of B-cell depletion further enhanced this ability, showing the enhanced effect of STING agonist and B-cell depletion in the control of HCC metastasis (**Fig. S6E**). This effect of STING agonism is remarkable, as none of the anti-VEGF-based combinations or anti-PD1 treatments have shown anti-metastatic effects in clinical studies or preclinical models of HCC.

### Combining STING agonism, ICB, and B-cell depletion eradicates tumor growth and prevents relapse in the RIL-175 HCC model

Since B-cell depletion overcame resistance in both STING agonist-treated and ICB-treated HCA-1 models individually, we next investigated its impact in a more potent setting combining STING agonism with dual PD1/VEGFR2 blockade (**Fig. 4A**). Strikingly, tumor growth was completely eradicated in this model, with all mice receiving the full combination treatment (STING agonist + anti-PD1/anti-VEGFR2 + anti-CD19/anti-B220) achieving complete responses (**Fig. 4B, C**). Moreover, rechallenging long-term survivors by implanting RIL-175 HCC cells into a different liver lobe resulted in no tumor growth, whereas tumors developed in age-matched control mice (**Fig. 4D, E**). These findings suggest that B-cell infiltration counteracts the therapeutic efficacy of STING agonism and dual PD-1/VEGFR2 blockade, potentially through immunosuppressive mechanisms. Notably, B-cell depletion significantly enhances tumor regression and leads to complete responses, indicating its role in overcoming resistance. Furthermore, the absence of tumor regrowth upon rechallenge highlights the establishment of durable anti-HCC immunological memory in this model.

**Figure 4.**
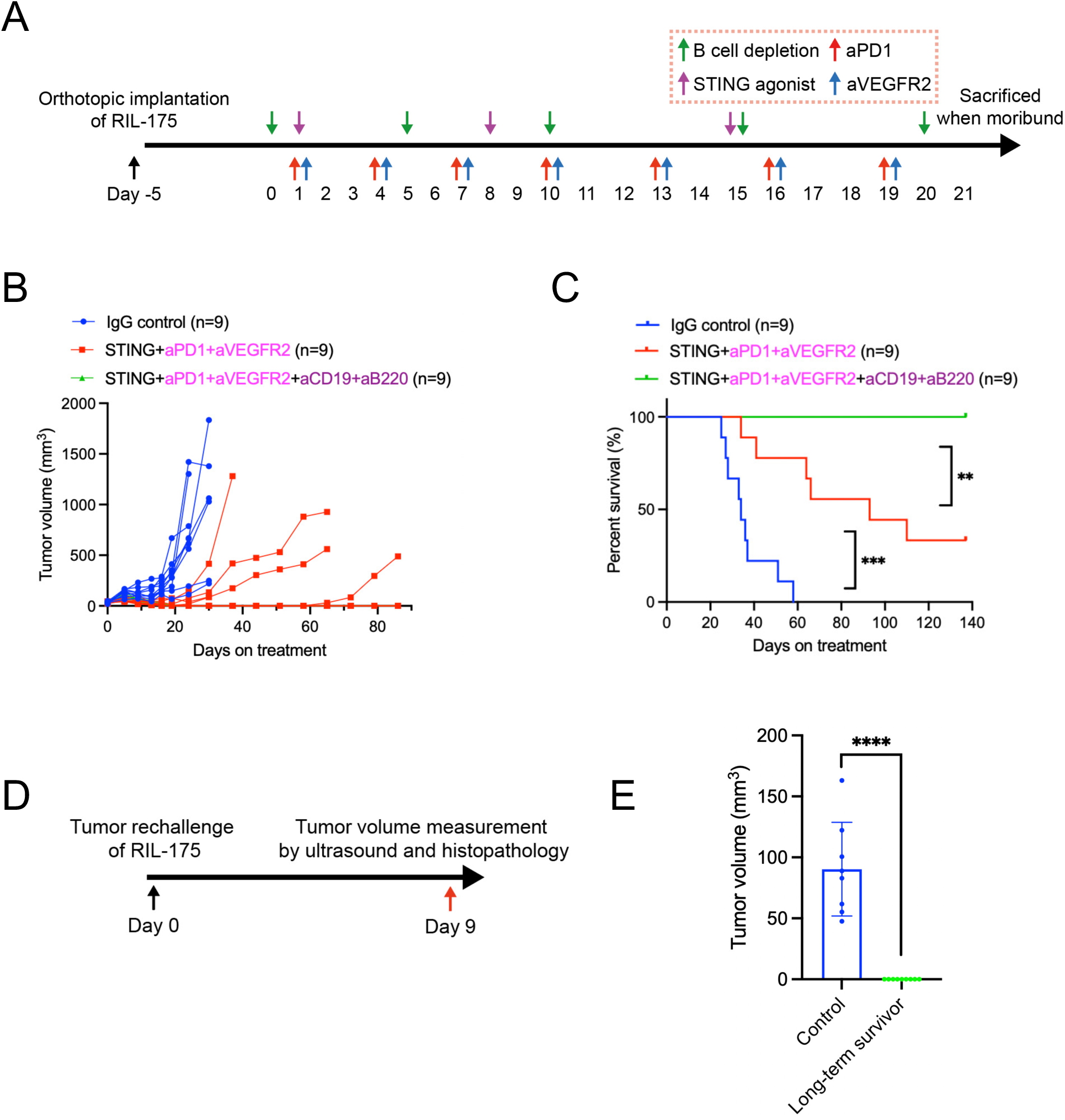
B cell depletion enhances STING agonism with dual PD1/VEGFR2 blockade in the RIL-175 murine HCC model. (**A**) Experimental design of B cell depletion and STING agonism with dual PD1/VEGFR2 blockade. (**B**) Tumor growth kinetics after treatment: the combination of STING agonist, dual anti-PD1/anti-VEGFR2, and B cell depletion group achieved complete tumor response in all mice, demonstrating a significantly superior tumor growth delay than other groups. (**C**) Overall survival of RIL-175 murine HCC-bearing mice after treatment: STING agonist with dual anti-PD1/anti-VEGFR2 plus B cell depletion induced a significant survival benefit than without B cell depletion or control groups. **, P < 0.01; ***, P < 0.001 from log-rank test, n=9/group. (**D**) Schematic for tumor rechallenge to the mice that have survived long-term from the initial survival cohort. (**E**) Comparison of tumor volumes detected by ultrasound 9 days after tumor rechallenge in long-term survivors (n=9) and age-matched control mice (n=7). ****, P < 0.0001 from the Mann-Whitney U test.

### TIM-1 expression is upregulated after STING agonist treatment in HCA-1 murine HCC

Given that B-cell infiltration in HCCs following immunotherapy may contribute to resistance, we analyzed the expression profiles of genes associated with B-cell regulatory functions using bulk RNA-sequencing data from HCC tissues in control and STING agonist-treated mice. This analysis revealed increased expression of regulatory Breg markers, including *Havcr1*, *Il10*, *Il12a*, and *Ebi3*, in HCA-1 HCC samples from STING agonist-treated mice (**Fig. 5A**). Based on these markers, we computed a “Breg score” using ssGSEA method (47), which was significantly higher in the tumor from STING agonist-treated group (**Fig. 5B**). Notably, *Havcr1*, which encodes the immune checkpoint TIM-1, was also upregulated in STING agonist-treated tumors compared to controls (**Fig. 5C**), suggesting an enrichment of TIM-1⁺ B cells following treatment. Furthermore, STING agonist-treated mice exhibited significantly higher plasma IL-10 levels, which may support the immunosuppressive role of infiltrating B cells in this context (**Fig. 5D**). In contrast, the same group displayed lower plasma levels of IFN-γ, IL-1β, and IL-2, indicating a reduction in pro-inflammatory cytokines (**Fig. S7A**).

**Figure 5.**
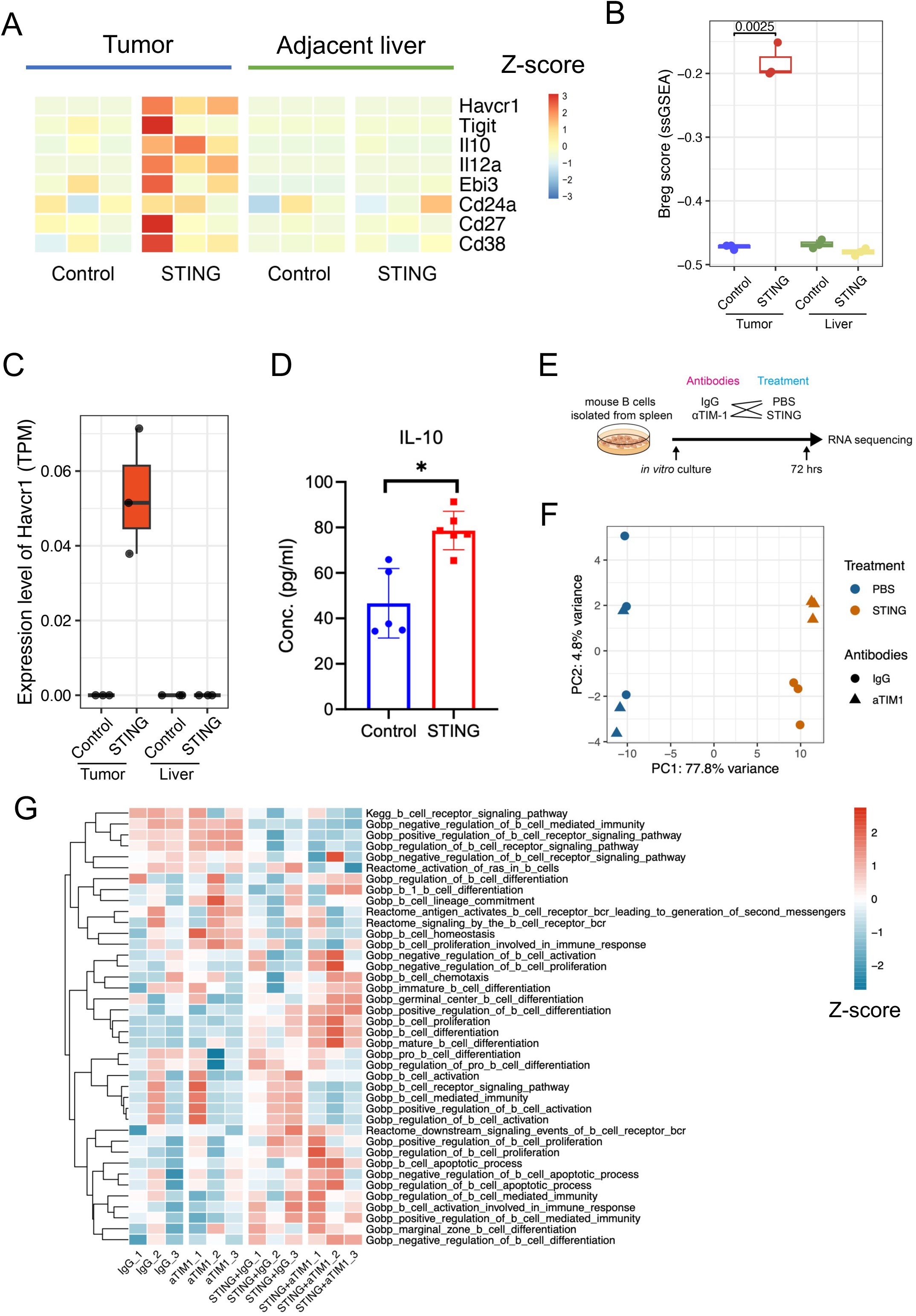
Higher TIM-1 expression in HCA-1 murine HCC after STING agonist treatment than control. (**A**) Heatmap showing the expression levels of Breg markers in both tumor and liver tissues from the control and STING agonist-treated groups. The color indicates the Z-score of gene expression levels. (**B**) Breg score comparison among tumor and liver tissues from the control and STING agonist-treated groups. (**C**) The expression level of the *Havcr1* gene was higher in the STING agonist-treated group. TPM, transcripts per million. (**D**) The concentration of IL-10 measured by ELISA was higher in the plasma collected from the STING agonist-treated group than in the control. *, P < 0.05, statistical significance was calculated by the Mann-Whitney U test. (**E**) Schematic for the in vitro treatment of B cells. (**F**) PCA plot showing the B cells clustered differently among groups. (**G**) The heatmap of the z-scores of different B-cell-related pathways demonstrates that STING agonism with anti-TIM-1 treatment induces B cell differentiation and proliferation.

To further investigate the role of STING agonism and TIM-1 signaling in B cells, we isolated B cells from mouse spleens and treated them in vitro with IgG, STING agonist, anti-TIM-1 blocking antibodies, or their combination. After 72 hours, RNA was extracted for transcriptomic analysis (**Fig. 5E**). The PCA showed a distinct transcriptional shift in STING agonist-treated B cells compared to the PBS control, with anti-TIM-1 treatment further enhancing this shift (**Fig. 5F**). Pathway enrichment analysis revealed that the combination of STING agonism and TIM-1 blockade promoted B-cell differentiation, including germinal center B-cell formation (**Fig. 5G**), and enhanced antigen processing and presentation (**Fig. S8**). These findings demonstrate that this combination therapy reprograms B cells, potentially altering their function within the TME.

### A combination of STING agonism and TIM-1 ICB is effective in an ICB-resistant murine HCC model

We conducted a survival experiment to test the efficacy of TIM-1 blockade in the context of STING agonism in the orthotopic anti-PD1-resistant HCA-1 model. Tumor-bearing mice received 15 days of treatment with (**i**) IgG control, (**ii**) STING agonist with IgG, (**iii**) anti-TIM-1 antibody, or (**iv**) STING agonist with anti-TIM-1 antibody (**Fig. S9A**). We found a significantly longer growth delay and median OS in the STING agonist/anti-TIM-1 combination group (**Fig. 6A, B**). Lung metastatic burden was also lowest in the combination group, in line with the anti-metastatic activity of STING agonism with B-cell targeting (**Figs. 6C, D** and **S9B**). Thus, combining STING agonism with TIM-1 blockade is a potentially effective treatment approach in anti-PD1-resistant HCC.

**Figure 6.**
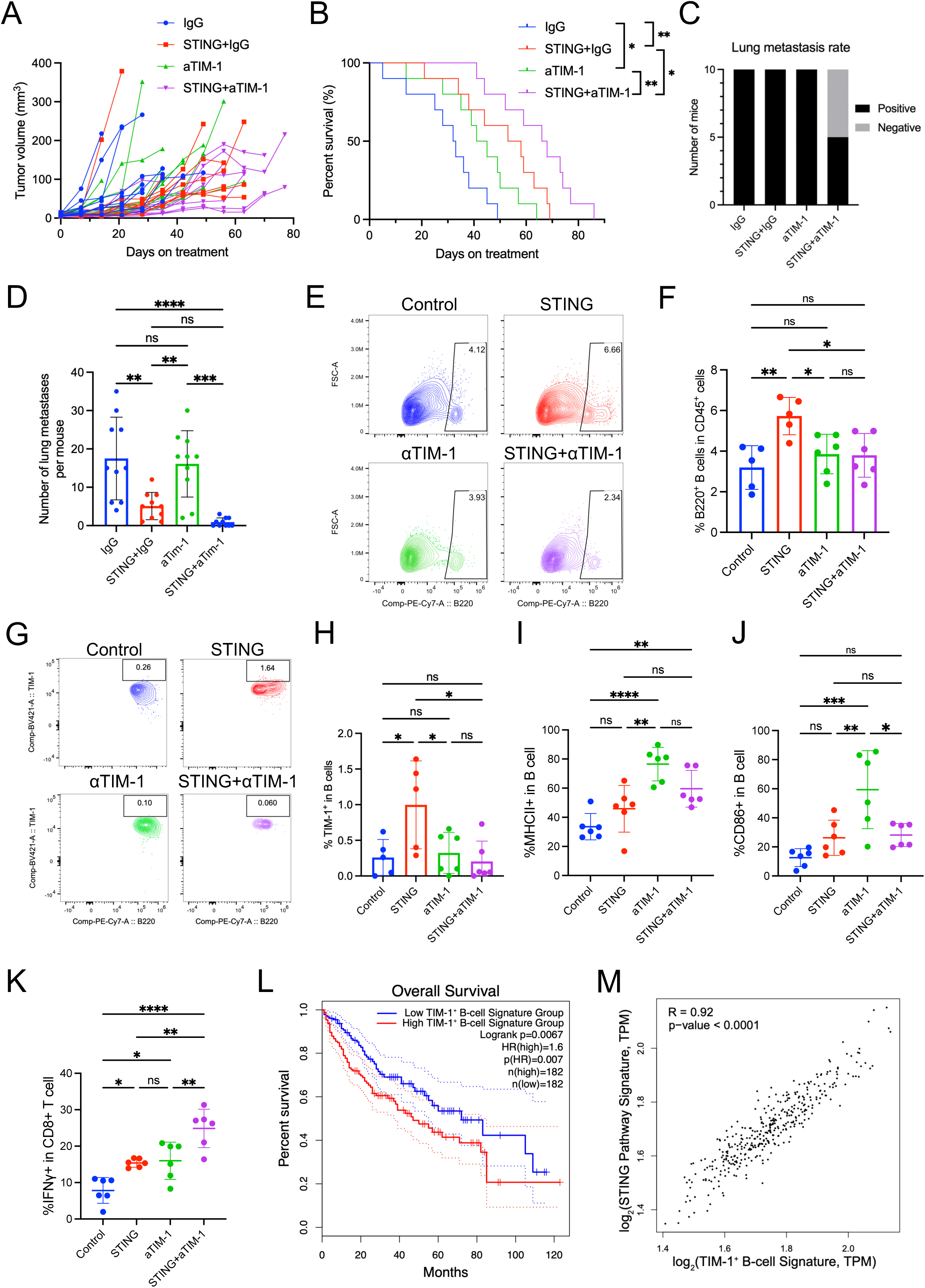
Combination treatment of STING agonism and TIM-1 blockade reprograms B cell function, induces more IFNγ^+^CD8^+^ T cells, and improves anti-tumor benefit. (**A**) Tumor growth kinetics after treatment: combined STING agonism and TIM-1 blockade induced tumor growth delay superior to the other groups. (**B**) Overall survival of HCC-bearing mice after treatment: STING agonist plus TIM-1 blockade induced a significant survival benefit compared to other groups. *, P < 0.05; **, P < 0.01 from log-rank test, n=10/group. (**C**) STING agonism combined with TIM-1 blockade reduced lung metastasis rates. (**D**) STING agonism/TIM-1 blockade combined treatment significantly reduced lung metastasis in the HCA-1 murine HCC model. P values were calculated by one-way ANOVA with the Tukey multiple comparisons test. (**E**, **F**) Flow cytometry analysis demonstrates increased B cell infiltration in the STING alone group. (**G**, **H**) Flow cytometry analysis demonstrates increased TIM-1^+^ B cells in the STING alone group, while the combination of TIM-1 blockade decreased its proportion. (**I**, **J**) The STING and TIM-1 blockade combination increased the proportion of MHCII^+^ and CD86^+^ B cells. (**K**) The STING and TIM-1 blockade combination increased the proportion of IFNγ^+^CD8^+^ T cells. *, P < 0.05; **, P < 0.01; ***, P < 0.001; ****, P < 0.0001, statistical significance was calculated by one-way ANOVA with Tukey’s test, n=6/group. (**L**) The prognostic value of the TIM-1^+^ B-cell signature in TCGA HCC patients (n=364, log-rank test). (**M**) The expression levels of TIM-1^+^ B-cell signature and STING pathway signature were positively correlated in TCGA HCC patients.

Next, to investigate how the combination of a STING agonist and anti-TIM-1 antibody reprograms the immune TME of HCC, we conducted a time-matched cohort study using the HCA-1 model in mice with liver damage. Mice with established tumors were randomized to treatment with (**i**) IgG control, (**ii**) STING agonist with IgG, (**iii**) anti-TIM-1 antibody, and (**iv**) STING agonist with anti-TIM-1 antibody. Tumor samples were collected on day 35 post-treatment to ensure sufficient material for flow cytometry analysis. Flow cytometry revealed that the STING agonist alone significantly increased B-cell infiltration while adding TIM-1 blockade reduced the B-cell frequency (**Fig. 6E, F**). TIM-1^+^ B cells were significantly elevated in the STING agonist group, and the addition of TIM-1 blockade prevented this increase (**Fig. 6G, H**). Among B cells, the proportions of MHC-II^+^ and CD86^+^ B cells were higher in groups treated with the anti-TIM-1 antibody (**Fig. 6I, J**), indicating an enhanced antigen-presenting capacity. Functionally, the combination treatment led to a significant increase in IFNγ^+^CD8^+^ T cells (**Fig. 6K**), further supporting the anti-tumor immune reprogramming of the TME of HCC induced by this therapeutic approach.

### TIM-1 and a B-cell signature are associated with survival in human HCC

To examine whether the expression levels of TIM-1^+^ B-cell signature (271 genes) correlate with survival in HCC patients, we used these genes to stratify the HCC patients from The Cancer Genome Atlas (TCGA) cohort into two groups of high and low scores based on the median. We found that this signature was significantly associated with the overall survival of these patients (**Fig. 6L**), and that the expression level of the *HAVCR1* gene alone was significantly associated with survival (**Fig. S9C**). Moreover, we found that the expression levels of TIM-1^+^ B-cell signature and STING pathway signature (54 genes) were directly correlated (**Fig. 6M**), further supporting the mechanistic link between the STING pathway activation and TIM-1^+^ B cells identified in HCC models.

To further explore the role of B cells in clinical HCC patients treated with ICB, we analyzed a single-cell RNA-seq dataset from a neoadjuvant clinical trial of ICB in HCC (59). Naive and memory B cells were examined from patients classified as anti-PD1 responders or non-responders (**Fig. S10A, B**). Differential gene expression analysis revealed that pathways associated with immune responses, including adaptive immune response, immune effector processes, and antigen processing and presentation, were enriched among the upregulated genes in responders (**Fig. S10C-E**). Among the significant differences in immune responses between responder and non-responder HCC patients, the scores for antigen processing and presentation, as well as B-cell activation, were significantly higher in the responder group (**Fig. S10F, G**), suggesting that functional activation of B cells is a critical target for ICB treatment efficacy.

## Discussion

ICB immunotherapy with PD1/PD-L1 blockers has transformed HCC treatment. However, inherent or acquired immunotherapy resistance remains a significant challenge, and the mechanisms driving resistance remain poorly understood. We studied resistance to ICB and STING agonism-based immunotherapy in well-characterized murine HCC models, which revealed a surprising mechanism of treatment resistance mediated by infiltrating B cells. We also show that targeting B-cell-mediated immunosuppression in HCC can prevent the acquired resistance to ICB and STING agonist therapy in murine HCC and identify the immune checkpoint TIM-1 as a novel target for HCC treatment (**Fig. 7**). As an immune checkpoint molecule on immunosuppressive B-cells, TIM-1 holds promise for guiding the development of new combination therapies that enhance the efficacy of STING agonists, offering new hope for the immunotherapy of HCC.

**Figure 7.**
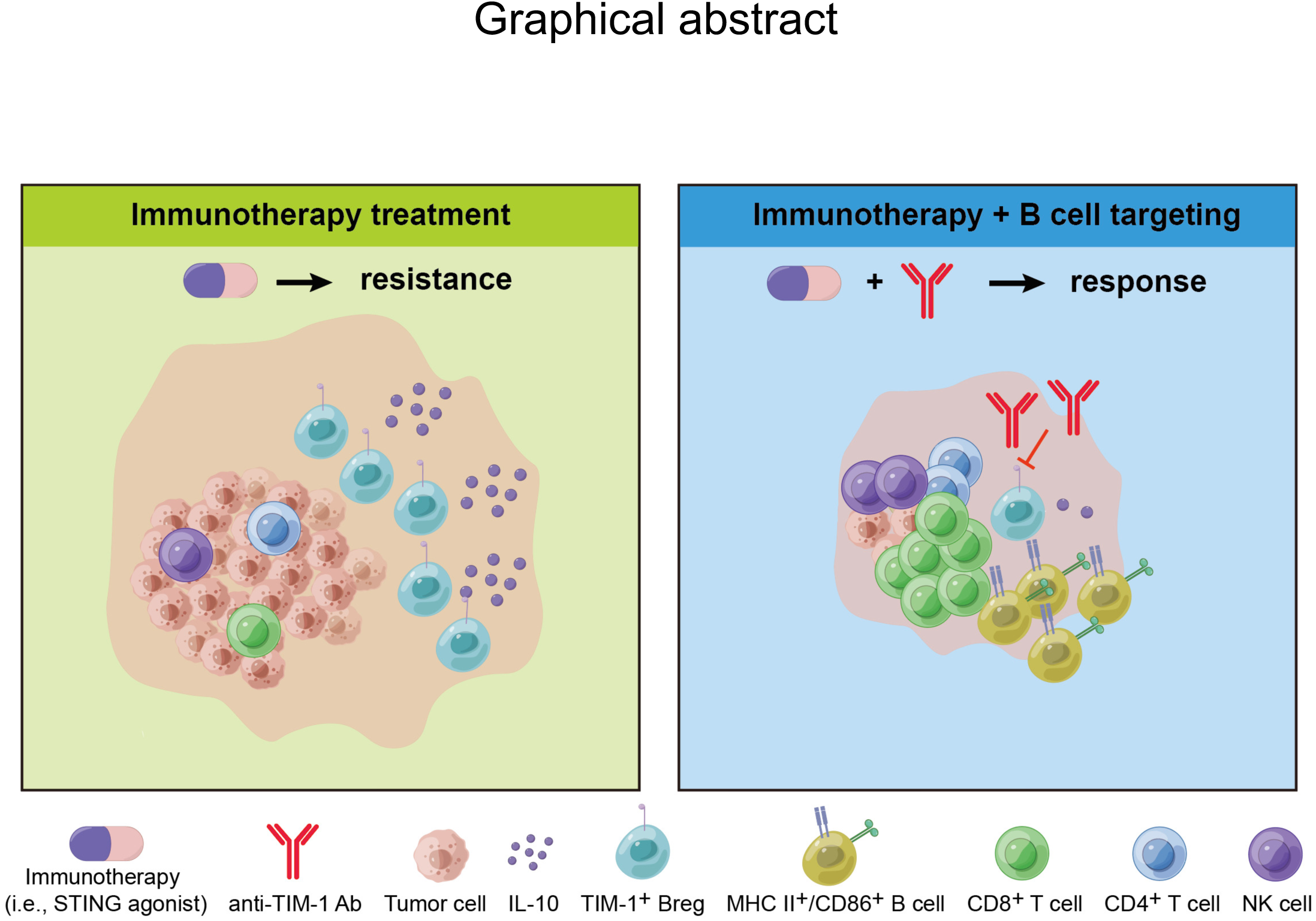
Targeting B cells enhances STING agonism in HCC. Immunotherapy resistance is partly mediated by IL-10-secreting TIM-1^+^ B cells in murine HCC. Combining immunotherapy with B-cell targeting therapy shows efficacy in murine HCC.

STING agonism showed potent activation of anti-tumor immunity but rapidly increased the infiltration by B-cells in the murine HCC tumor tissue, particularly in the immunotherapy-resistant HCA-1 model. Considering the pleiotropic roles of B-cells in cancer progression, we used the depletion of B-cells to clarify their predominant role in this treatment setting. The overall depletion of B-cells significantly enhanced the efficacy of STING agonism, including when combined with STING agonist and anti-PD1 therapy, demonstrating the predominantly immunosuppressive role of B-cell populations during immunotherapy in HCC. This discovery points to B-cells as potential targets for overcoming immunotherapy resistance. Indeed, combining STING agonism, dual anti-PD1/VEGFR2 therapy, and B cell depletion led to HCC eradication in all mice and immunological memory responses. Approaches to specifically target immunosuppressive B-cells in HCC still need to be developed. We show here that combining ICB using antibodies blocking TIM-1, an immune checkpoint expressed on immunosuppressive and tumor-promoting B-cells (22), significantly improves survival in ICB-resistant murine HCC.

Lung metastasis is the most common form of distant dissemination in liver cancer patients and is associated with treatment resistance and poor outcomes (55–57). A recent study showed that STING activity increases in metastatic progenitors re-entering the cell cycle (60). In contrast, another study showed that persistent activation of STING leads to desensitization and rewiring of downstream signaling, which impedes effective anti-tumor immunity and may instead facilitate cancer metastasis (61). We found that combining a STING agonist with B-cell depletion significantly reduced the incidence of lung metastases, lung metastatic burden, and pleural effusion. Moreover, combining anti-TIM-1 ICB treatment with a STING agonist also inhibited lung metastasis, offering a new strategy to improve outcomes in metastatic HCC.

Of note, we found an increased formation of B-cell aggregates and TLS-like structures in HCC tissues from STING agonist-treated mice. The function of the B cells in the TLS-like structures in HCC may explain the conflicting data on the association between TLSs and outcomes in cancer patients, as our results show that B-cells promoted immunosuppression after immunotherapy. The balance between positive and negative feedback from different T cell subsets is believed to govern the functionality of tumor-associated TLS, with Bregs potentially contributing to additional negative feedback (62). Interestingly, IL-10-producing Bregs in TLS-like aggregates in breast cancer patients are associated with shorter metastasis-free survival (63). Further studies in HCC are needed to dissect the pro-versus anti-tumor roles of TLS to better understand their impact on tumor progression and therapeutic response. Given the lack of Breg-specific markers (64), our finding of TIM-1 as targeting to prevent or convert the Breg phenotype may represent a more promising approach than depletion.

Agonists activating the STING pathway have shown modest anti-cancer efficacy in clinical trials so far (28,38,39), and ICB-based therapy is ineffective in most HCC patients. Despite potently activating anti-tumor immunity, resistance to STING agonists and ICB occurs frequently and rapidly, mediated by immunosuppressive mechanisms. Our study showed that B cells are a major source of therapy resistance under both innate and acquired immune response activated by STING agonist and ICB treatment, and identified that combining STING agonist with B-cell targeting and anti-TIM-1 ICB can improve the treatment response in murine HCC. These strategies should be tested clinically to treat patients with HCC.

Overall, our findings reveal a critical role for B cells in shaping the immune TME and influencing immunotherapy outcomes in HCC. While STING agonists and ICB alone have shown limited efficacy, targeting B-cell-mediated immunosuppression could enhance anti-tumor immunity. More broadly, our study underscores the role of B cells in hindering immunotherapy responses. By dissecting these mechanisms and functional states of B cells, we provide a foundation for new combination strategies to reprogram the TME and improve the efficacy of ICB, STING agonism, and other immunotherapies in HCC.

## Funding Information

This study was supported by a sponsored research agreement with Bristol Myers Squibb (BMS). DGD’s research is supported by NIH grants R01CA260857, R01CA254351, R01CA247441, R03CA256764, and P01CA261669, and Department of Defense PRCRP grants W81XWH-19-1-0284 and W81XWH-21-1-0738. L.B was supported by grants from the Cancer Research Institute (CRI), the Bridge project, the Lung Cancer Research Foundation (LCRF) and the MGH Transformative Scholars Program.

## Author disclosures

DGD reports research grants from Bayer, Exelixis, and Surface Oncology, outside the submitted work. The other authors reported no disclosures.

## Authors’ Contributions

Conceptualization: XL, ZL, DGD

Methodology: XL, ZL, TK, PL, YS, DY, JW, ML, AM, KM, PH, MK, LB

Investigation: XL, ZL, TK, PL, YS, DY, JW, ML, AM, KM

Visualization: XL, ZL, TK, AM

Funding acquisition: DGD

Project administration: DGD

Supervision: DGD

Writing – original draft: XL, DGD

Writing – review & editing: all authors

## Acknowledgments

The authors sincerely thank M. Duquette, A. Khachatryan, C. Smith, H. Taniguchi, and S. Roberge (all MGH) for outstanding technical support.

## References

1. Llovet JM, Castet F, Heikenwalder M, Maini MK, Mazzaferro V, Pinato DJ, et al. Immunotherapies for hepatocellular carcinoma. Nat Rev Clin Oncol 2022;19:151–72.

2. Villanueva A. Hepatocellular Carcinoma. N Engl J Med 2019;380:1450–62.

3. Vogel A, Meyer T, Sapisochin G, Salem R, Saborowski A. Hepatocellular carcinoma. Lancet 2022;400:1345–62.

4. Llovet JM, Kelley RK, Villanueva A, Singal AG, Pikarsky E, Roayaie S, et al. Hepatocellular carcinoma. Nat Rev Dis Primers 2021;7:6.

5. Rumgay H, Arnold M, Ferlay J, Lesi O, Cabasag CJ, Vignat J, et al. Global burden of primary liver cancer in 2020 and predictions to 2040. J Hepatol 2022;77:1598–606.

6. Llovet JM, Montal R, Sia D, Finn RS. Molecular therapies and precision medicine for hepatocellular carcinoma. Nat Rev Clin Oncol 2018;15:599–616.

7. Yang X, Yang C, Zhang S, Geng H, Zhu AX, Bernards R, et al. Precision treatment in advanced hepatocellular carcinoma. Cancer Cell 2024;42:180–97.

8. Abou-Alfa GK, Lau G, Kudo M, Chan SL, Kelley RK, Furuse J, et al. Tremelimumab plus Durvalumab in Unresectable Hepatocellular Carcinoma. NEJM Evid 2022;1:EVIDoa2100070.

9. Finn RS, Qin S, Ikeda M, Galle PR, Ducreux M, Kim TY, et al. Atezolizumab plus Bevacizumab in Unresectable Hepatocellular Carcinoma. N Engl J Med 2020;382:1894–905.

10. Pardoll DM. The blockade of immune checkpoints in cancer immunotherapy. Nat Rev Cancer 2012;12:252–64.

11. Sharonov GV, Serebrovskaya EO, Yuzhakova DV, Britanova OV, Chudakov DM. B cells, plasma cells and antibody repertoires in the tumour microenvironment. Nat Rev Immunol 2020;20:294–307.

12. Cyster JG, Allen CDC. B Cell Responses: Cell Interaction Dynamics and Decisions. Cell 2019;177:524–40.

13. Downs-Canner SM, Meier J, Vincent BG, Serody JS. B Cell Function in the Tumor Microenvironment. Annu Rev Immunol 2022;40:169–93.

14. Cabrita R, Lauss M, Sanna A, Donia M, Skaarup Larsen M, Mitra S, et al. Tertiary lymphoid structures improve immunotherapy and survival in melanoma. Nature 2020;577:561–5.

15. Calderaro J, Petitprez F, Becht E, Laurent A, Hirsch TZ, Rousseau B, et al. Intra-tumoral tertiary lymphoid structures are associated with a low risk of early recurrence of hepatocellular carcinoma. J Hepatol 2019;70:58–65.

16. Helmink BA, Reddy SM, Gao J, Zhang S, Basar R, Thakur R, et al. B cells and tertiary lymphoid structures promote immunotherapy response. Nature 2020;577:549–55.

17. Petitprez F, de Reynies A, Keung EZ, Chen TW, Sun CM, Calderaro J, et al. B cells are associated with survival and immunotherapy response in sarcoma. Nature 2020;577:556–60.

18. Michaud D, Steward CR, Mirlekar B, Pylayeva-Gupta Y. Regulatory B cells in cancer. Immunol Rev 2021;299:74–92.

19. Xiao X, Lao XM, Chen MM, Liu RX, Wei Y, Ouyang FZ, et al. PD-1hi Identifies a Novel Regulatory B-cell Population in Human Hepatoma That Promotes Disease Progression. Cancer Discov 2016;6:546–59.

20. Shalapour S, Font-Burgada J, Di Caro G, Zhong Z, Sanchez-Lopez E, Dhar D, et al. Immunosuppressive plasma cells impede T-cell-dependent immunogenic chemotherapy. Nature 2015;521:94–8.

21. Shalapour S, Lin XJ, Bastian IN, Brain J, Burt AD, Aksenov AA, et al. Inflammation-induced IgA+ cells dismantle anti-liver cancer immunity. Nature 2017;551:340–5.

22. Bod L, Kye YC, Shi J, Torlai Triglia E, Schnell A, Fessler J, et al. B-cell-specific checkpoint molecules that regulate anti-tumour immunity. Nature 2023;619:348–56.

23. Ablasser A, Chen ZJ. cGAS in action: Expanding roles in immunity and inflammation. Science 2019;363.

24. Kong X, Zuo H, Huang HD, Zhang Q, Chen J, He C, et al. STING as an emerging therapeutic target for drug discovery: Perspectives from the global patent landscape. J Adv Res 2023;44:119–33.

25. Ishikawa H, Barber GN. STING is an endoplasmic reticulum adaptor that facilitates innate immune signalling. Nature 2008;455:674–8.

26. Sun L, Wu J, Du F, Chen X, Chen ZJ. Cyclic GMP-AMP synthase is a cytosolic DNA sensor that activates the type I interferon pathway. Science 2013;339:786–91.

27. Wu J, Sun L, Chen X, Du F, Shi H, Chen C, et al. Cyclic GMP-AMP is an endogenous second messenger in innate immune signaling by cytosolic DNA. Science 2013;339:826–30.

28. Huang C, Shao N, Huang Y, Chen J, Wang D, Hu G, et al. Overcoming challenges in the delivery of STING agonists for cancer immunotherapy: A comprehensive review of strategies and future perspectives. Mater Today Bio 2023;23:100839.

29. Li T, Chen ZJ. The cGAS-cGAMP-STING pathway connects DNA damage to inflammation, senescence, and cancer. J Exp Med 2018;215:1287–99.

30. Marcus A, Mao AJ, Lensink-Vasan M, Wang L, Vance RE, Raulet DH. Tumor-Derived cGAMP Triggers a STING-Mediated Interferon Response in Non-tumor Cells to Activate the NK Cell Response. Immunity 2018;49:754–63 e4.

31. Zhang C, Shang G, Gui X, Zhang X, Bai XC, Chen ZJ. Structural basis of STING binding with and phosphorylation by TBK1. Nature 2019;567:394–8.

32. Luke JJ, Piha-Paul SA, Medina T, Verschraegen CF, Varterasian M, Brennan AM, et al. Phase I Study of SYNB1891, an Engineered E. coli Nissle Strain Expressing STING Agonist, with and without Atezolizumab in Advanced Malignancies. Clin Cancer Res 2023;29:2435–44.

33. Meric-Bernstam F, Sweis RF, Hodi FS, Messersmith WA, Andtbacka RHI, Ingham M, et al. Phase I Dose-Escalation Trial of MIW815 (ADU-S100), an Intratumoral STING Agonist, in Patients with Advanced/Metastatic Solid Tumors or Lymphomas. Clin Cancer Res 2022;28:677–88.

34. Meric-Bernstam F, Sweis RF, Kasper S, Hamid O, Bhatia S, Dummer R, et al. Combination of the STING Agonist MIW815 (ADU-S100) and PD-1 Inhibitor Spartalizumab in Advanced/Metastatic Solid Tumors or Lymphomas: An Open-Label, Multicenter, Phase Ib Study. Clin Cancer Res 2023;29:110–21.

35. Li S, Mirlekar B, Johnson BM, Brickey WJ, Wrobel JA, Yang N, et al. STING-induced regulatory B cells compromise NK function in cancer immunity. Nature 2022;610:373–80.

36. Flood BA, Higgs EF, Li S, Luke JJ, Gajewski TF. STING pathway agonism as a cancer therapeutic. Immunol Rev 2019;290:24–38.

37. Galon J, Bruni D. Approaches to treat immune hot, altered and cold tumours with combination immunotherapies. Nat Rev Drug Discov 2019;18:197–218.

38. Amouzegar A, Chelvanambi M, Filderman JN, Storkus WJ, Luke JJ. STING Agonists as Cancer Therapeutics. Cancers (Basel) 2021;13.

39. Samson N, Ablasser A. The cGAS-STING pathway and cancer. Nat Cancer 2022;3:1452–63.

40. Jiang M, Chen P, Wang L, Li W, Chen B, Liu Y, et al. cGAS-STING, an important pathway in cancer immunotherapy. J Hematol Oncol 2020;13:81.

41. Zhou J, Sun H, Wang Z, Cong W, Wang J, Zeng M, et al. Guidelines for the Diagnosis and Treatment of Hepatocellular Carcinoma (2019 Edition). Liver Cancer 2020;9:682–720.

42. Tofilon PJ, Basic I, Milas L. Prediction of in vivo tumor response to chemotherapeutic agents by the in vitro sister chromatid exchange assay. Cancer Res 1985;45:2025–30.

43. Reiberger T, Chen Y, Ramjiawan RR, Hato T, Fan C, Samuel R, et al. An orthotopic mouse model of hepatocellular carcinoma with underlying liver cirrhosis. Nat Protoc 2015;10:1264–74.

44. Kapanadze T, Gamrekelashvili J, Ma C, Chan C, Zhao F, Hewitt S, et al. Regulation of accumulation and function of myeloid derived suppressor cells in different murine models of hepatocellular carcinoma. J Hepatol 2013;59:1007–13.

45. Bankhead P, Loughrey MB, Fernandez JA, Dombrowski Y, McArt DG, Dunne PD, et al. QuPath: Open source software for digital pathology image analysis. Sci Rep 2017;7:16878.

46. Aran D, Hu Z, Butte AJ. xCell: digitally portraying the tissue cellular heterogeneity landscape. Genome Biol 2017;18:220.

47. Barbie DA, Tamayo P, Boehm JS, Kim SY, Moody SE, Dunn IF, et al. Systematic RNA interference reveals that oncogenic KRAS-driven cancers require TBK1. Nature 2009;462:108–12.

48. Tang Z, Kang B, Li C, Chen T, Zhang Z. GEPIA2: an enhanced web server for large-scale expression profiling and interactive analysis. Nucleic Acids Res 2019;47:W556–W60.

49. Chen J, Amoozgar Z, Liu X, Aoki S, Liu Z, Shin SM, et al. Reprogramming the Intrahepatic Cholangiocarcinoma Immune Microenvironment by Chemotherapy and CTLA-4 Blockade Enhances Anti-PD-1 Therapy. Cancer Immunol Res 2024;12:400–12.

50. Morita S, Lei PJ, Shigeta K, Ando T, Kobayashi T, Kikuchi H, et al. Combination CXCR4 and PD1 blockade enhances intratumoral dendritic cell activation and immune responses against hepatocellular carcinoma. Cancer Immunol Res 2024.

51. Shigeta K, Datta M, Hato T, Kitahara S, Chen IX, Matsui A, et al. Dual Programmed Death Receptor-1 and Vascular Endothelial Growth Factor Receptor-2 Blockade Promotes Vascular Normalization and Enhances Antitumor Immune Responses in Hepatocellular Carcinoma. Hepatology 2020;71:1247–61.

52. Shigeta K, Matsui A, Kikuchi H, Klein S, Mamessier E, Chen IX, et al. Regorafenib combined with PD1 blockade increases CD8 T-cell infiltration by inducing CXCL10 expression in hepatocellular carcinoma. J Immunother Cancer 2020;8.

53. Xiao Y, Chen J, Zhou H, Zeng X, Ruan Z, Pu Z, et al. Combining p53 mRNA nanotherapy with immune checkpoint blockade reprograms the immune microenvironment for effective cancer therapy. Nat Commun 2022;13:758.

54. Gulen MF, Koch U, Haag SM, Schuler F, Apetoh L, Villunger A, et al. Signalling strength determines proapoptotic functions of STING. Nat Commun 2017;8:427.

55. Katyal S, Oliver JH, 3rd, Peterson MS, Ferris JV, Carr BS, Baron RL. Extrahepatic metastases of hepatocellular carcinoma. Radiology 2000;216:698–703.

56. Ou L, Lu G, Cao M, Hu M. Lung metastases after liver cancer resection cured by immunotherapy: case report and literature review. Anticancer Drugs 2023;34:e1–e8.

57. Wu C, Ren X, Zhang Q. Incidence, risk factors, and prognosis in patients with primary hepatocellular carcinoma and lung metastasis: a population-based study. Cancer Manag Res 2019;11:2759–68.

58. Chen Y, Huang Y, Reiberger T, Duyverman AM, Huang P, Samuel R, et al. Differential effects of sorafenib on liver versus tumor fibrosis mediated by stromal-derived factor 1 alpha/C-X-C receptor type 4 axis and myeloid differentiation antigen-positive myeloid cell infiltration in mice. Hepatology 2014;59:1435–47.

59. Magen A, Hamon P, Fiaschi N, Soong BY, Park MD, Mattiuz R, et al. Intratumoral dendritic cell-CD4(+) T helper cell niches enable CD8(+) T cell differentiation following PD-1 blockade in hepatocellular carcinoma. Nat Med 2023;29:1389–99.

60. Hu J, Sanchez-Rivera FJ, Wang Z, Johnson GN, Ho YJ, Ganesh K, et al. STING inhibits the reactivation of dormant metastasis in lung adenocarcinoma. Nature 2023;616:806–13.

61. Li J, Hubisz MJ, Earlie EM, Duran MA, Hong C, Varela AA, et al. Non-cell-autonomous cancer progression from chromosomal instability. Nature 2023;620:1080–8.

62. Laumont CM, Nelson BH. B cells in the tumor microenvironment: Multi-faceted organizers, regulators, and effectors of anti-tumor immunity. Cancer Cell 2023;41:466–89.

63. Ishigami E, Sakakibara M, Sakakibara J, Masuda T, Fujimoto H, Hayama S, et al. Coexistence of regulatory B cells and regulatory T cells in tumor-infiltrating lymphocyte aggregates is a prognostic factor in patients with breast cancer. Breast Cancer 2019;26:180–9.

64. Bodogai M, Lee Chang C, Wejksza K, Lai J, Merino M, Wersto RP, et al. Anti-CD20 antibody promotes cancer escape via enrichment of tumor-evoked regulatory B cells expressing low levels of CD20 and CD137L. Cancer Res 2013;73:2127–38.

