## Supplementary Figures for "Inhibiting B-cell-mediated Immunosuppression to Enhance the Immunotherapy Efficacy in Hepatocellular Carcinoma"

### Figure S1

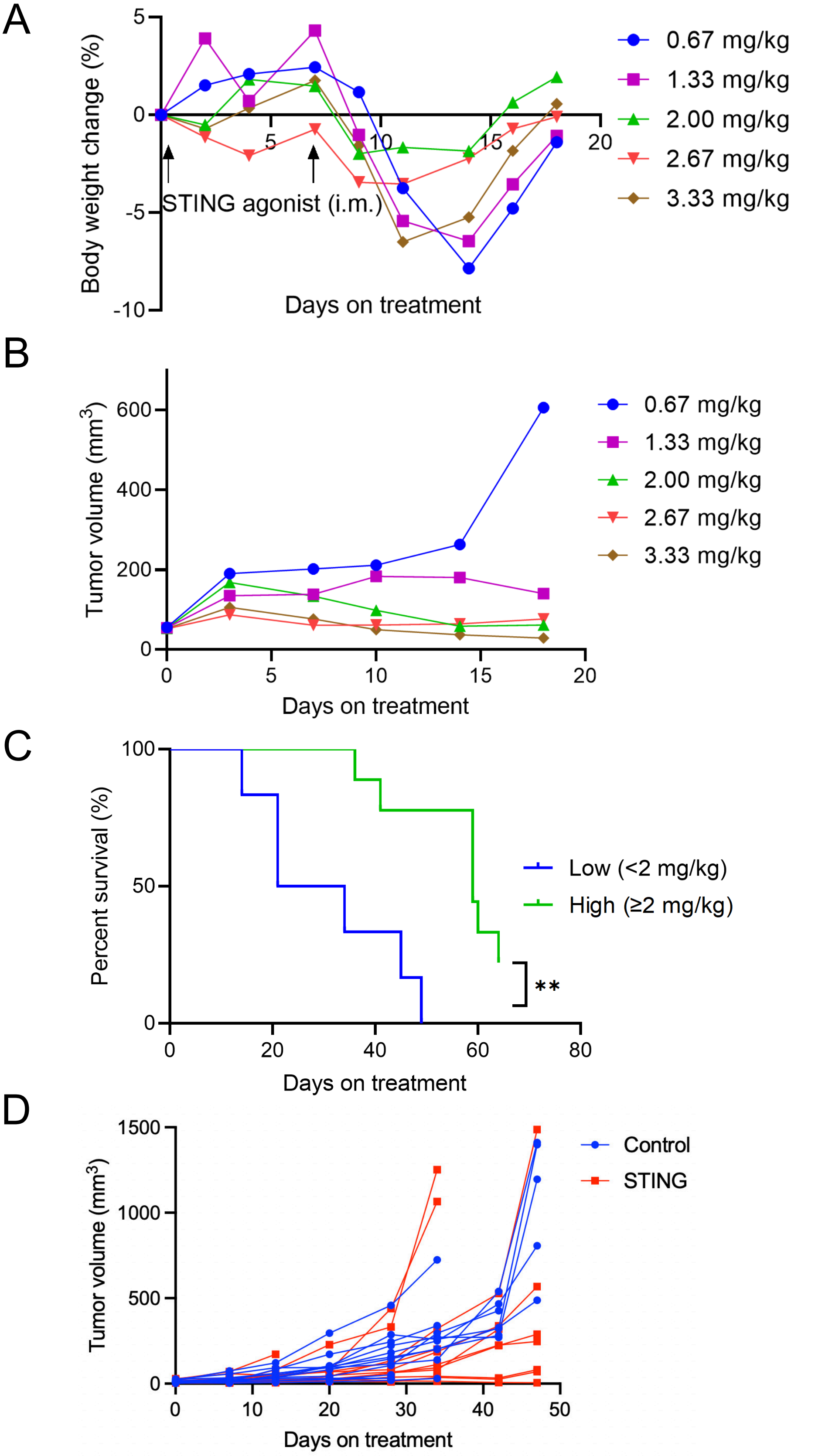


**Figure S1: Safety and feasibility of STING agonist treatment *in vivo*.** (**A**) Mean body weight changes after treatment with STING agonist under different doses. i.m., intramuscular injection. (**B**) Tumor growth kinetics after treatment with different doses of STING agonist in the orthotopic RIL-175 murine HCC model with liver damage. Data are shown as mean tumor volumes. (**C**) Overall survival of HCC-bearing mice after treatment. A low concentration was defined as less than 2 mg/kg, and a high concentration was defined as more than or equal to 2 mg/kg. n=3 mice at each concentration. (**D**) Individual tumor growth kinetics after 2-dose treatment in orthotopic HCA-1 HCC model: STING agonist demonstrated a non-significant trend toward delayed tumor growth compared to the control group.

Figure S2

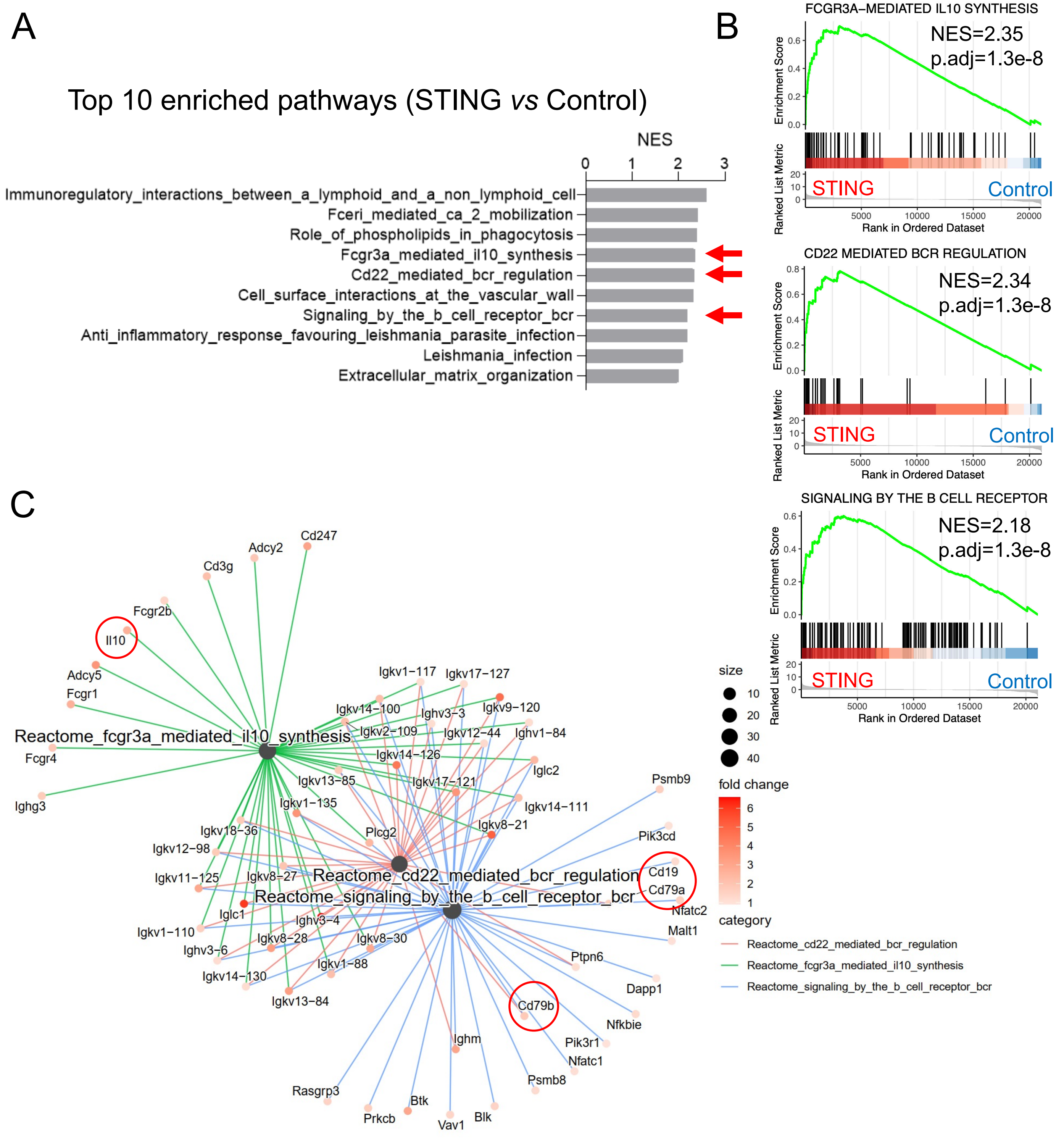


**Figure S2: Bulk RNA-seq analysis of tumor tissues from the STING agonist versus control treatment groups.** (**A**) Top 10 enriched pathway terms (Reactome database) in HCC samples in STING agonist-treated group compared with control. Three B-cell-related pathways are pointed with red arrows. (**B**) Enrichment of FCGR3A-mediated IL10 synthesis, CD22-mediated BCR regulation, and signaling by the B cell receptor pathways in STING agonist-treated group versus control group by gene set enrichment analysis (GSEA). NES, normalized enrichment score; p.adj, adjusted p-value. (**C**) Network plot showing the genes of the enriched B-cell-related pathways.

### Figure S3

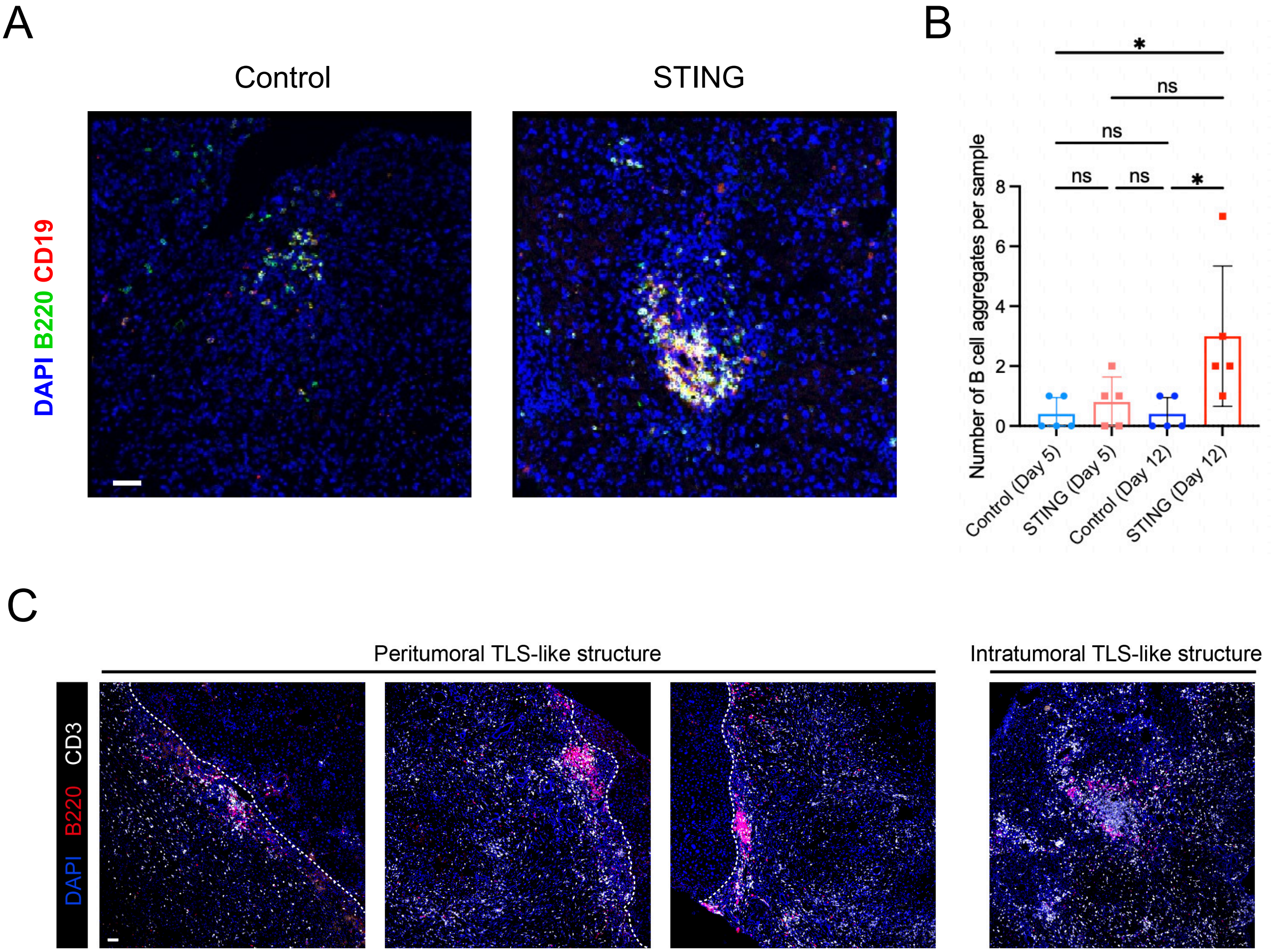


**Figure S3: Increased formation of B cell aggregates after STING agonist treatment in the RIL-175 HCC murine model.** (**A**) Representative IF results for the B cell aggregates found in RIL-175 HCC tumors (scale bar, 50μm). (**B**) STING agonist treatment induced more B cell aggregates than the control group on day 12 after the treatment. (**C**) Representative IF images of peritumoral and intratumoral TLS-like structures. The boundary between the tumor and the adjacent liver was shown as a dotted line.

Figure S4

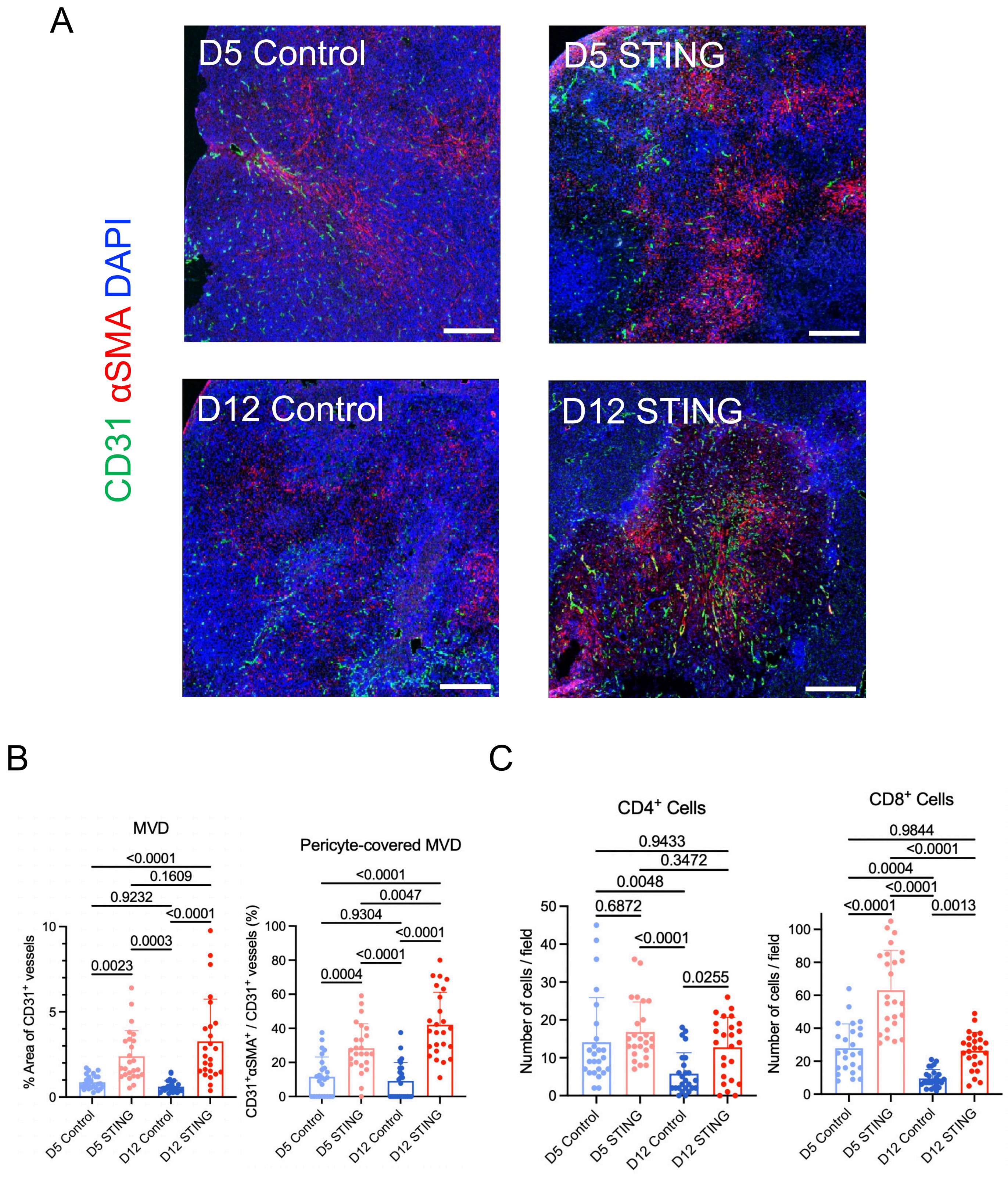


**Figure S4: TME changes in RIL-175 murine HCC model after STING agonist treatment.** (**A**) Representative immunofluorescence images of tumor vessels in RIL-175 HCC tumor tissues, collected on days 5 and 12 after initiating treatment. Scale bar, 500μm. (**B**) Total and pericyte-covered microvessel density (MVD) in the four treatment groups. (**C**) Frequency of $CD4^+$ and $CD8^+$ tumor-infiltrating T cells in the four treatment groups. P values were calculated by one-way ANOVA with Tukey multiple comparisons test.

### Figure S5

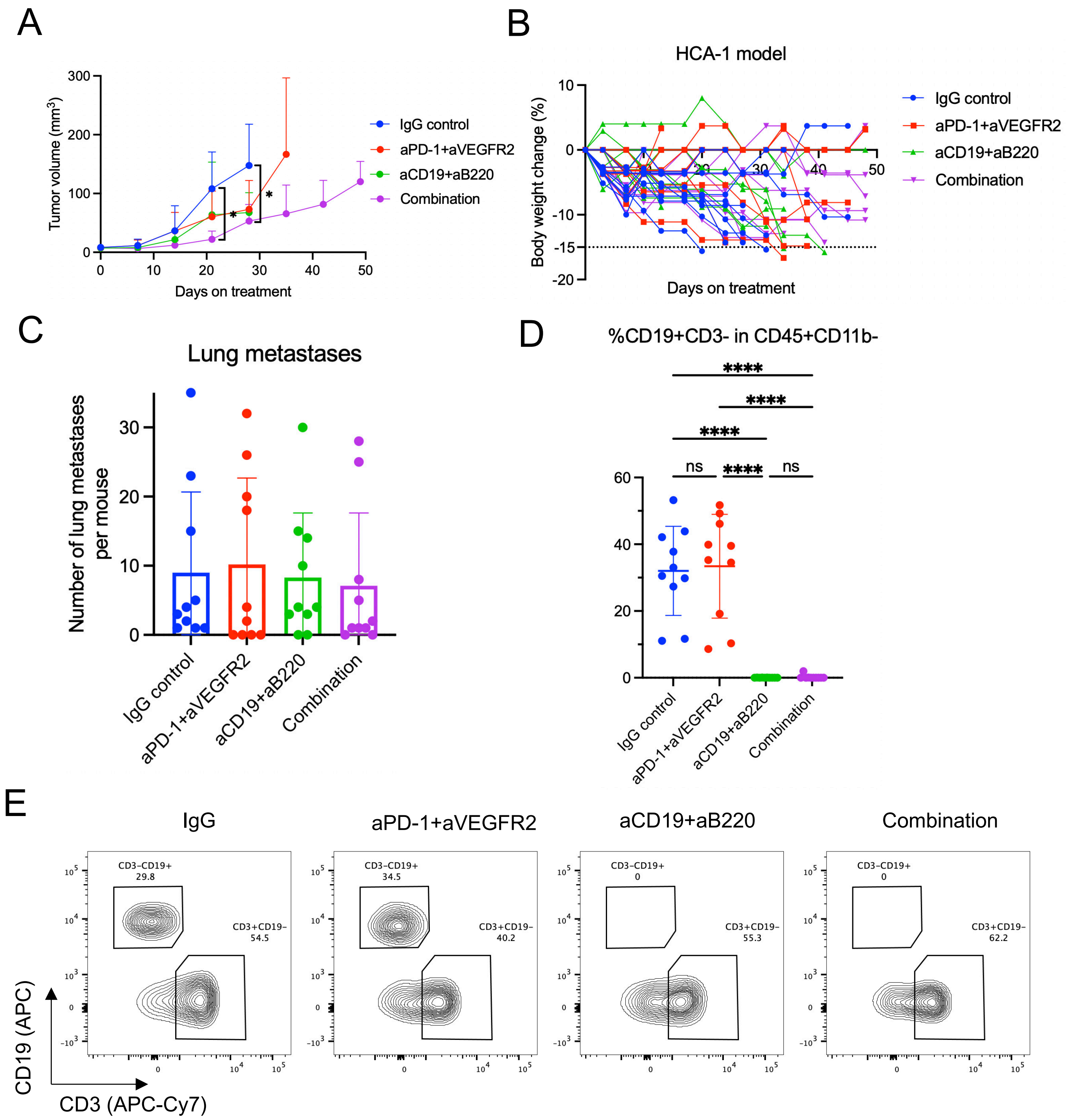


**Figure S5: Combination treatment of dual anti-PD1/VEGFR2 blockade and B cell depletion in the HCA-1 murine HCC model.** (**A**) Individual tumor growth curves after treatment: the combination of dual anti-PD1/VEGFR2 blockade and B cell depletion group induced tumor growth delay superior to other groups. (**B**) Individual body weight changes after treatment in the HCA-1 murine model. (**C**) No significant differences were found in lung metastases among the groups. (**D**) The proportions of $CD19^{+}CD3^{-}$ B cells in $CD45^{+}CD11b^{-}$ cells are significantly lower in B cell depletion alone group and combination group. (**E**) Representative flow cytometry analysis demonstrates effective B cell depletion in the B cell depletion alone and the combination groups. n=10 mice/group.

### Figure S6

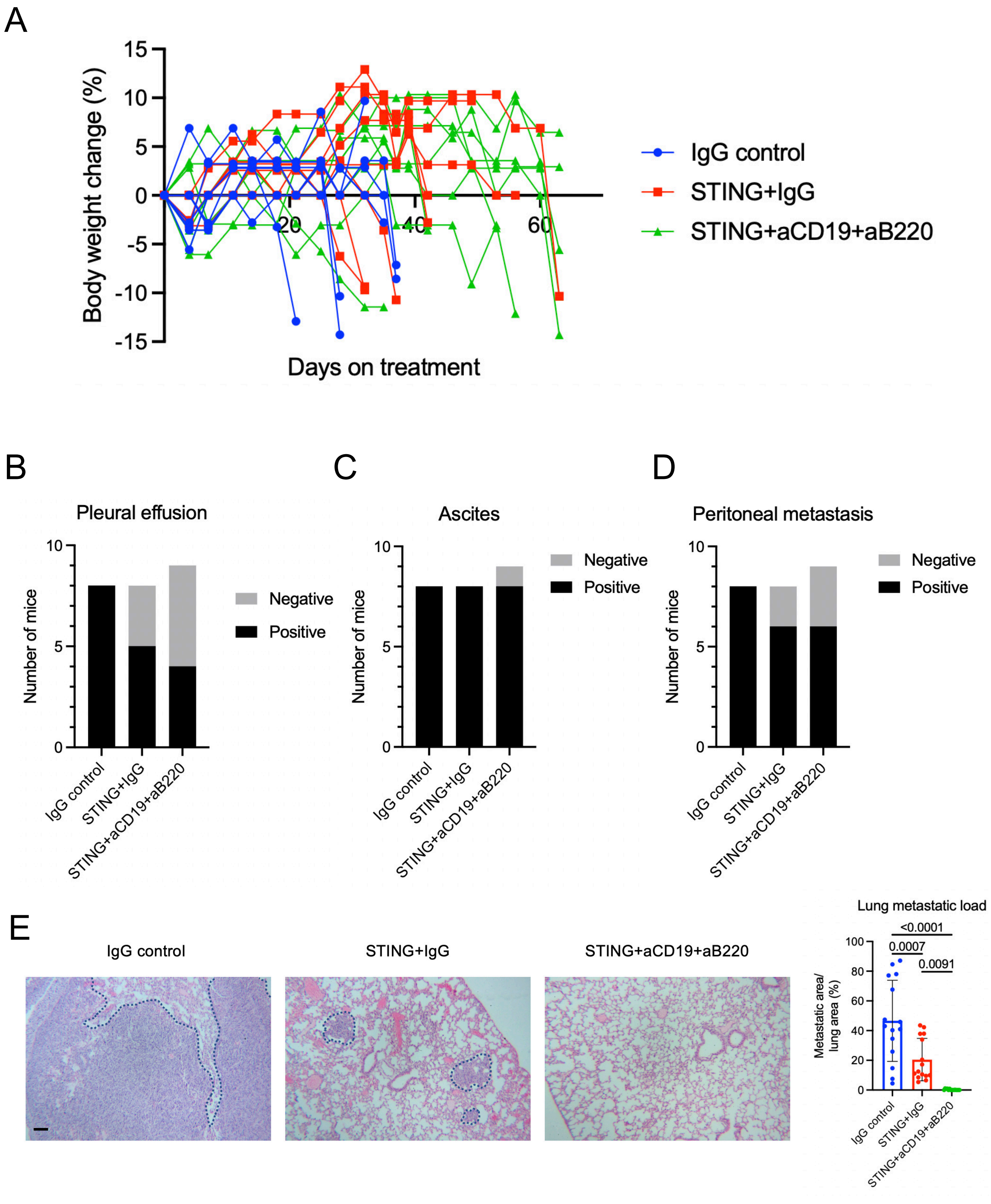


**Figure S6: Combination of B-cell depletion and STING agonism in the HCA-1 murine HCC model.** (**A**) Individual body weight changes after treatment in each group. (**B-D**) Pleural effusion (**B**), ascites (**C**), and peritoneal metastasis (**D**) incidence in each group. (**E**) H&E staining showing lung metastasis in the HCA-1 murine HCC model (scale bar, 200μm). Areas representing tumors are delineated with dotted lines. STING agonism/B cell depletion combination treatment shows significantly lower lung metastatic load as the percentage of metastatic area in the lung area. P values were calculated by one-way ANOVA with the Tukey multiple comparisons test.

### Figure S7

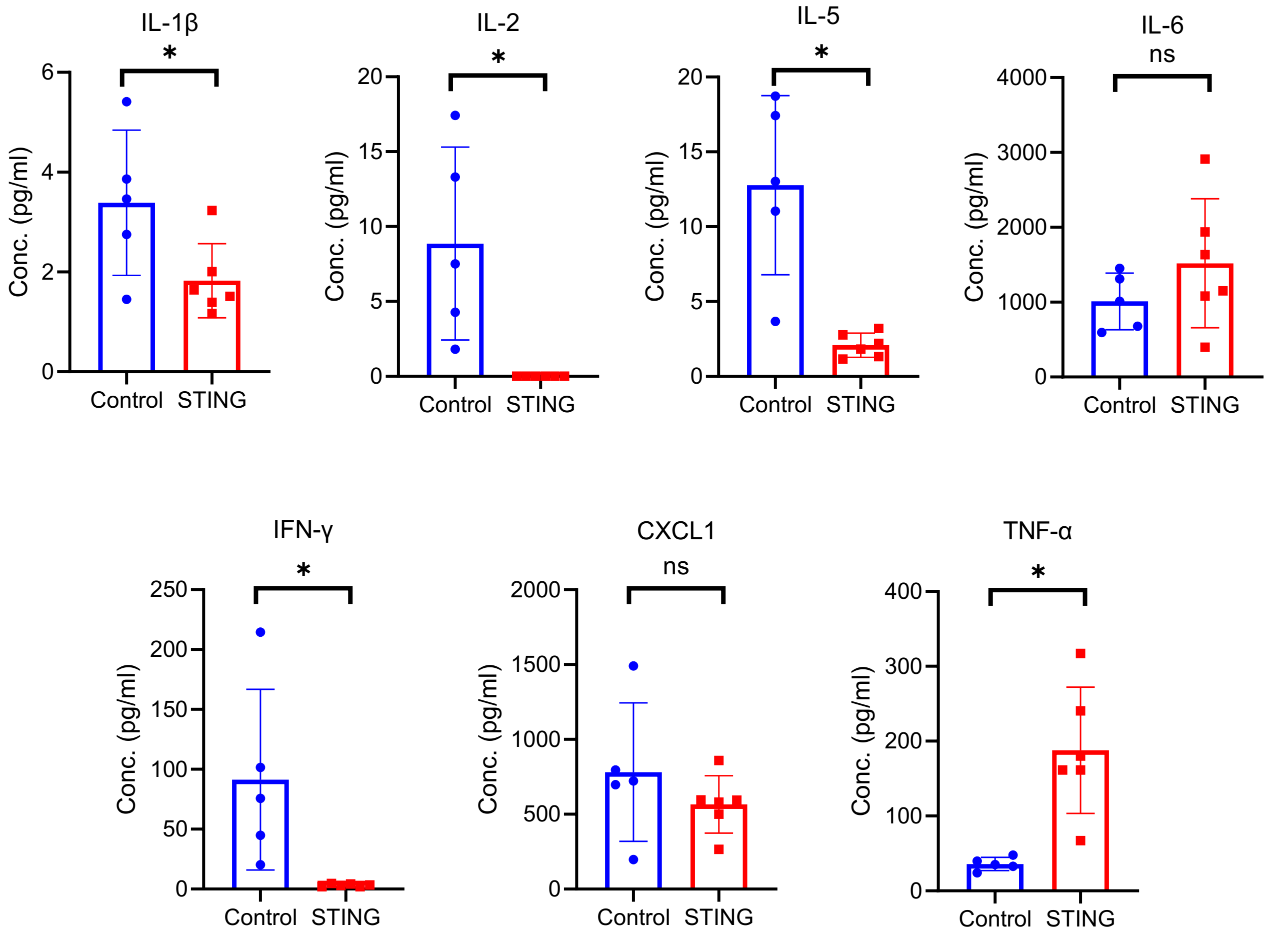


**Figure S7: Multiplex array analysis of cytokine and chemokine changes after STING agonist treatment in plasma samples from HCA-1 murine HCC-bearing mice.** Plasma samples were collected on day 20 after starting treatment. P values were calculated using an unpaired t-test.

### Figure S8

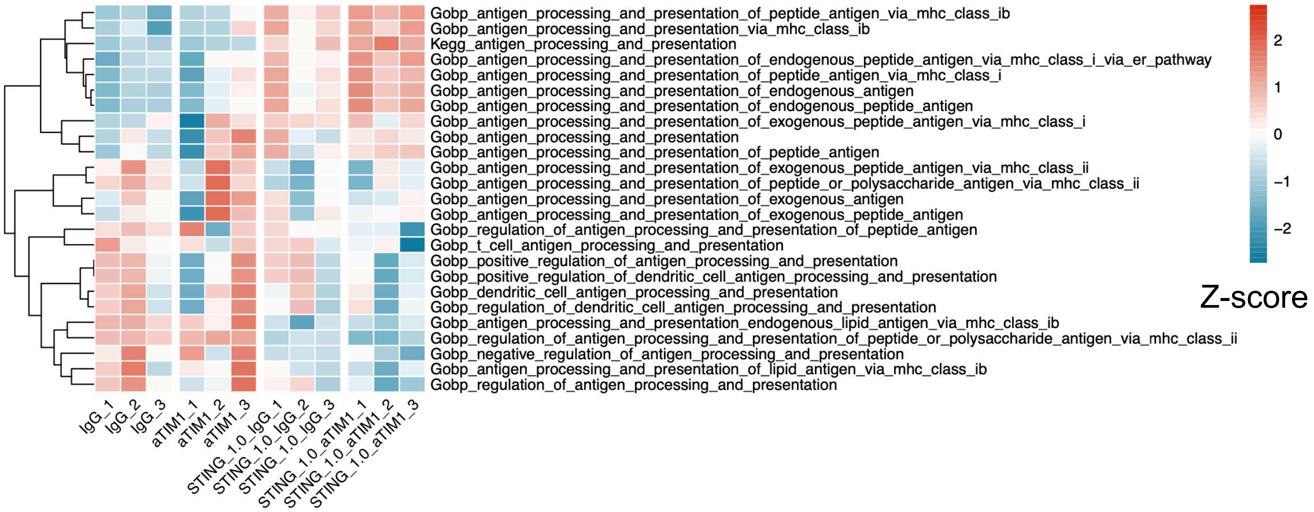


**Figure S8: Antigen processing and presentation pathway enrichment analysis in B cells cultured *in vitro*.** The heatmap of the z-score of different antigen-related pathways demonstrates that STING agonism combined with anti-TIM-1 treatment induces more antigen processing and presentation in GO-BP and KEGG pathway databases.

### Figure S9

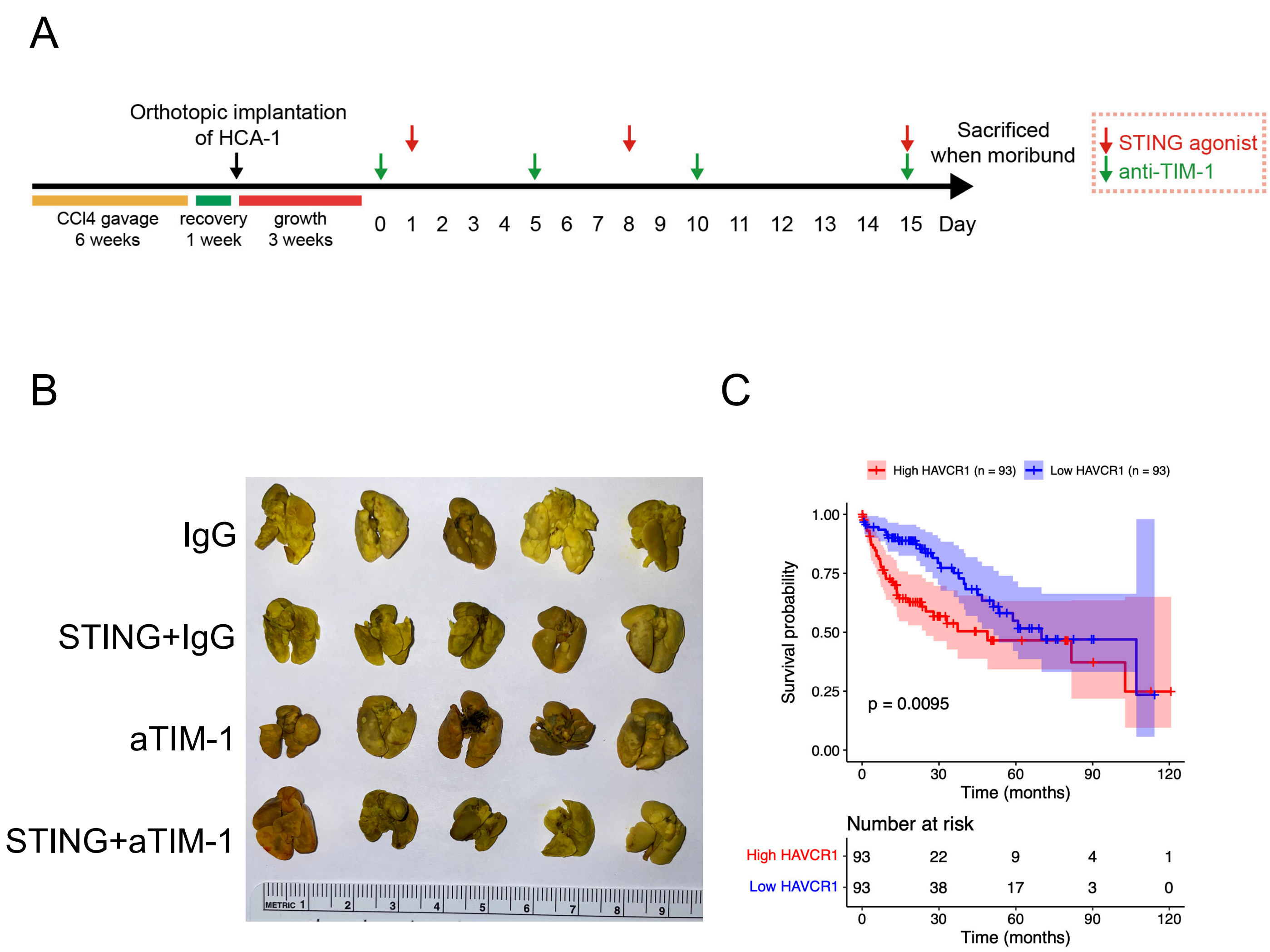


**Figure S9: TIM-1 as a targeting in HCA-1 model treatment.** (**A**) Experimental design of TIM-1 blockade with STING agonist treatment. (**B**) Representative photographs of lungs after fixation in Bouin's solution. (**C**) The prognostic value of the *HAVCR1* gene in TCGA HCC patients. Upper quartile vs lower quartiles; P values were calculated by log-rank test.

Figure S10

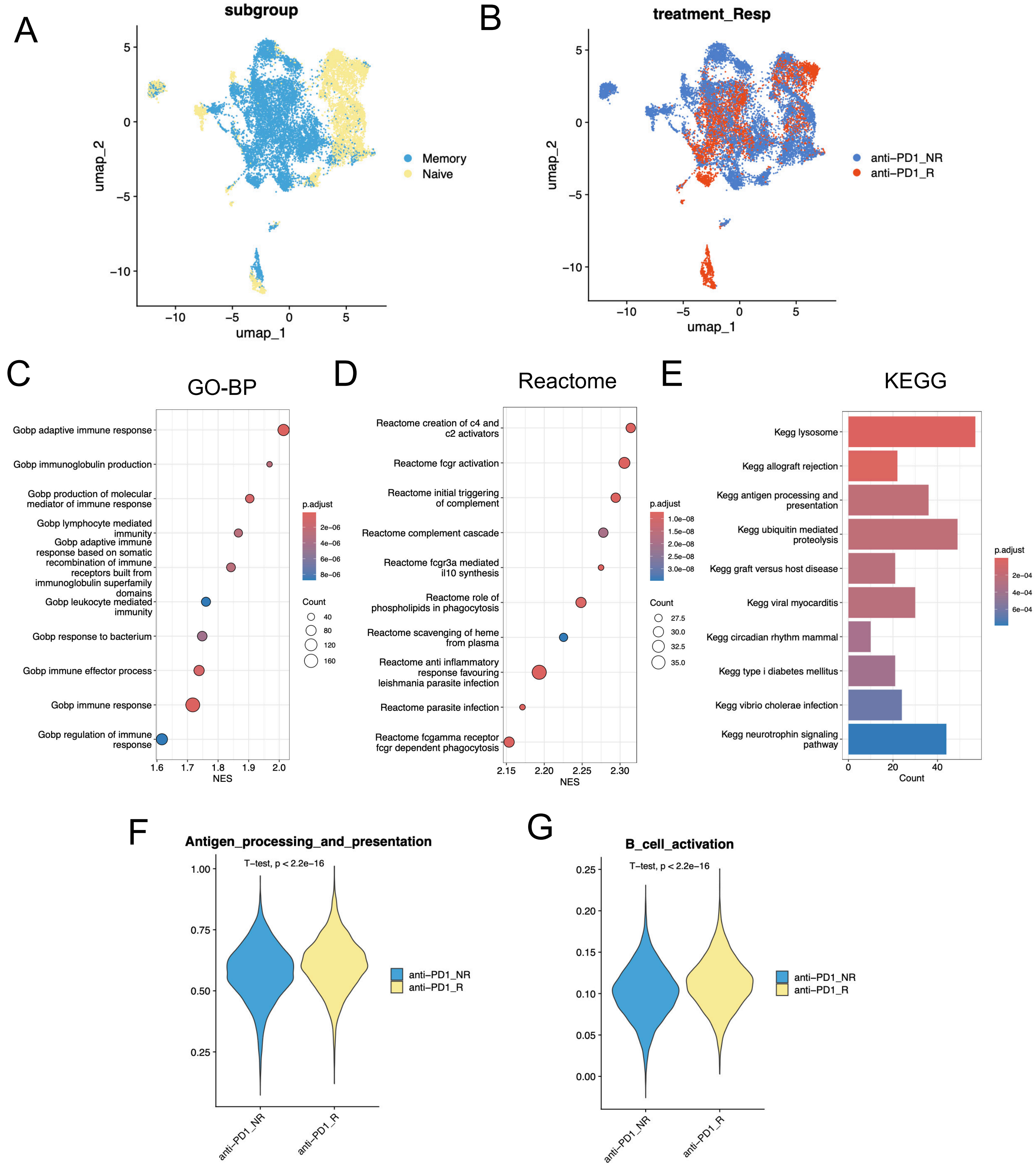


**Figure S10: Analysis of publically available single-cell RNA-seq dataset from tumor samples from human HCC patients treated with neoadjuvant anti-PD-1 treatment.** (**A**) The u-map plot demonstrates the memory and naive B cell subtypes. (**B**) The u-map plot demonstrates the B cells from patients labeled as non-response or response. (**C**, **D**) GSEA analysis of up-regulated genes in responders vs non-responders in GO-BP (**C**) and Reactome (**D**) databases. (**E**) Pathway enrichment analysis of up-regulated genes in responders vs non-responders in the KEGG database. (**F**, **G**) Scores of antigen processing and presentation (**F**) and B cell activation (**G**) were increased in anti-PD-1 responders than in non-responders. P values were calculated by t-test.
